# McSplicer: a probabilistic model for estimating splice site usage from RNA-seq data

**DOI:** 10.1101/2020.08.10.243097

**Authors:** Israa Alqassem, Yash Sonthalia, Erika Klitzke-Feser, Heejung Shim, Stefan Canzar

## Abstract

Alternative splicing removes intronic sequences from transcripts in alternative ways to produce different forms (isoforms) of mature mRNA. The composition of expressed transcripts and their alternative forms give specific functionalities to cells in a particular condition or developmental stage. In addition, a large fraction of human disease mutations affect splicing and lead to aberrant mRNA and protein products. Current methods that interrogate the transcriptome based on RNA-seq either suffer from short read length when trying to infer full-length transcripts, or are restricted to predefined units of alternative splicing that they quantify from local read evidence. Instead of attempting to quantify individual outcomes of the splicing process such as local splicing events or full-length transcripts, we propose to quantify alternative splicing using a simplified probabilistic model of the underlying splicing process. Our model is based on the usage of individual splice sites and can generate arbitrarily complex types of splicing patterns. In our method, McSplicer, we estimate the parameters of our model using all read data at once and we demonstrate in our experiments that this yields more accurate estimates compared to competing methods. Our model is able to describe multiple effects of splicing mutations using few, easy to interpret parameters, as we illustrate in an experiment on RNA-seq data from autism spectrum disorder patients. McSplicer is implemented in Python and available as open-source at https://github.com/canzarlab/McSplicer.

## 1 Introduction

Through alternative splicing (AS), a single gene can produce multiple mRNA transcripts, or isoforms, that combine exons in alternative ways. Approximately 95% of human multi-exon protein-coding genes undergo alternative splicing [34], creating a remarkably complex set of transcripts that give specific functionalities to cells and tissues in a particular condition or developmental stage. Even more than in healthy tissue, aberrant splicing is prevalent in various diseases, including cancer and neurologic diseases [13, 51].

RNA sequencing (RNA-seq) is routinely used in genome-wide transcript analysis. This technology produces short reads from which existing methods infer and quantify RNA splicing, broadly, in one of two different ways. Methods either analyze full-length transcripts or focus on individual splicing events. Transcript assembly methods such as StringTie [37], CIDANE [7], and CLASS [45] aim to identify the set of expressed full-length transcripts which in principle provides a complete picture of all splicing variations, see e.g. transcript *t*_1_-*t*_5_ in Fig. 1. Since the number of possible transcripts that can be stitched together from short reads is larger than the number of truly expressed transcripts and each read carries little information about its originating transcript, the transcript assembly problem is ill-posed [26] and error-prone especially for complex genes expressing multiple transcript isoforms [18, 1].

**Figure 1:**
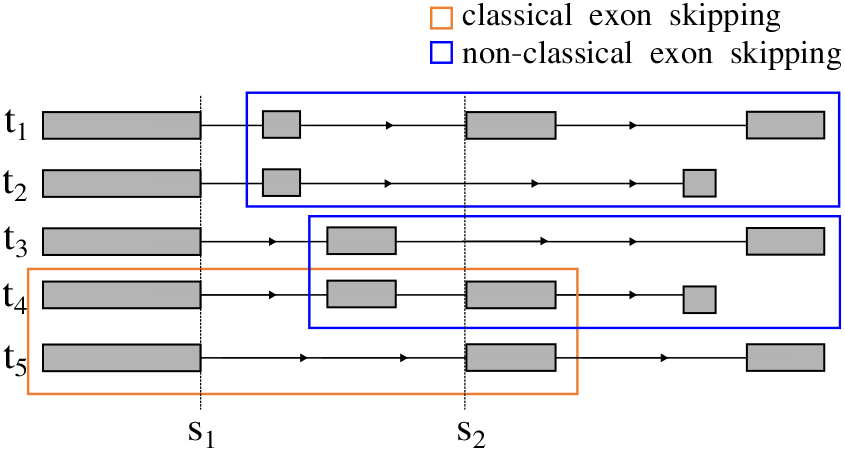
Complex alternative splicing involving 5 different transcripts. The two classical exon skipping events between *t*_1_ and *t*_5_, and between *t*_4_ and *t*_5_ do not fully capture the overall complexity. The two exon skippings marked in blue are not considered classical events and would not be reported by methods such as SplAdder, since they also differ in the last exon. Methods such as MAJIQ generalize simple events to more complex AS units that contain all introns sharing a common splice site. Two such AS units are required to describe the simple exon skipping event marked in orange, one comprising three introns sharing donor *s*_1_ and one containing three different introns sharing acceptor *s*_2_. In contrast, McSplicer estimates the usage of individual splice sites. The usage of acceptor site *s*_2_, for example, would be lowered by both exon skipping events marked in blue.

Event-based methods, therefore, focus on local splicing patterns such as the classical exon skipping event denoted in Fig. 1, without a prior attempt to assemble full-length transcripts (with the notable exception of SUPPA [2]). The relative abundance of different splicing outcomes that can potentially be shared by multiple transcripts, can then be quantified using a simple metric such as percent spliced in (PSI) [50]. Event-based methods differ in the complexity of the units of AS they quantify. In the simplest case, methods such as MISO [24], SUPPA, ASGAL [9], SpliceGrapher [40], and SplAdder [23] identify one of the canonical types of AS, such as exon skipping, alternative 5’ and 3’ splice sites, and intron retentions (see Supplementary Figure S1). These simple types of splicing events describe two possible splicing outcomes that differ in a single exon or splice site. In Fig. 1, this definition would include the two simple exon skippings between *t*_1_ and *t*_5_, and between *t*_4_ and *t*_5_, clearly underestimating the full AS complexity across *t*_1_-*t*_5_.

Complex events, on the other hand, involve multiple alternative splice sites or exons and according to [48] constitute at least one-third of AS events observed in human and mouse tissues. Methods such as JUM [52], MAJIQ [48] and the splice-site-centric quantification method proposed in [33], therefore consider AS units that generalize simple events to more complex patterns. They quantify the relative usage of an arbitrary number of introns that share a common splice site, such as the three introns sharing common donor site *s*_1_ in Fig. 1. Since these AS units capture only the common endpoints of alternative splicing patterns, such methods need to quantify two AS units for a single exon skipping event (Fig. 1). LeafCutter [30], on the other hand, iteratively extends AS units by introns that share a common splice site with any of the current set of introns. At the extreme end, Whippet [46] includes all introns spanning or connecting an alternative splicing event. It then enumerates all possible transcript fragments that combine overlapping events and estimates their relative abundance using an EM algorithm similar to full-length transcript quantification methods such as kallisto [5].

Here, we propose to quantify alternative splicing by building a probabilistic model as a simple approximation to the underlying splicing processes, rather than focusing on individual outcomes of the processes such as local splicing events or full-length transcripts. Previously, LeGault et al. [27] proposed a similar idea, quantifying alternative splicing using generative probabilistic models that weight edges of splicing graphs [19], using parameters representing RNA processing conditional probabilities. These parameters can then be used to estimate transcript and processing event frequencies. In this work, we propose a simpler probabilistic model. Our model employs the usages of annotated as well as novel splice sites across all expressed transcripts to describe a simplified splicing process that has generated the set of expressed transcripts. Traversing the linear ordering of all exons of a gene from 5’ to 3’, the usage of each splice site specifies the probability with which the site is used as donor or acceptor site. For example, the usage of acceptor *s*_2_ in Fig. 1 indicates the abundance of transcripts *t*_1_, *t*_4_, and *t*_5_ that “use” the acceptor relative to the total output *t*_1_-*t*_5_ of the gene. Our model assumes that splice site usages are independent of each other, leading to a simpler model compared to the model in [27].

This model by definition can generate complex splicing patterns that do not rely on any pre-defined simple or complex AS units as event-based methods like SplAdder, MAJIQ, or LeafCutter do. At the same time, splice site usages that capture simultaneous changes in multiple isoforms facilitate the interpretation of point mutations that disrupt splicing as is the case in many genetic disorders [3]. Instead of attempting to quantify each one of multiple possible effects on intron or even transcript level, a reduced splice site usage as computed by McSplicer may directly reflect the weakening of a splice site by a point mutation in the consensus splice site sequence that is responsible for these effects, as we illustrate in our experiments on RNA-seq data from autism spectrum disorder patients (Section 3.4).

Furthermore, our method simultaneously estimates the model parameters, i.e. splice site usages, using all reads mapped to a gene locus, often resulting in more accurate estimates compared to other methods, e.g. event-based methods, that use only reads directly supporting their parameters. We demonstrate the improved accuracy of McSplicer compared to existing methods in our experiments.

## 2 Method

A typical RNA-seq analysis workflow that uses McSplicer to estimate the usage of splice sites consist of the five steps illustrated in Fig. 2. After A) mapping reads in a RNA-seq sample to a reference genome sequence using a read alignment tool such as STAR [10] or HISAT [25], we B) assemble reads to full-length transcripts using methods such as StringTie [37] or CIDANE [7] to identify annotated as well as novel splice sites. We use the extracted splice sites to C) partition a gene into contiguous, non-overlapping segments. We count reads that overlap distinct combinations, or *signatures*, of such segments. Reads that map to the same signature are equivalent in terms of the splicing pattern they represent [7]. From *signature counts*, i.e. the number of reads mapping to different signatures (see Supplementary Figure S2 for an illustration), McSplicer estimates splice site usages in step (D). Splice site usages computed by McSplicer can be leveraged in E) different types of downstream anlyses, including the quantification of various types of splicing events.

**Figure 2:**
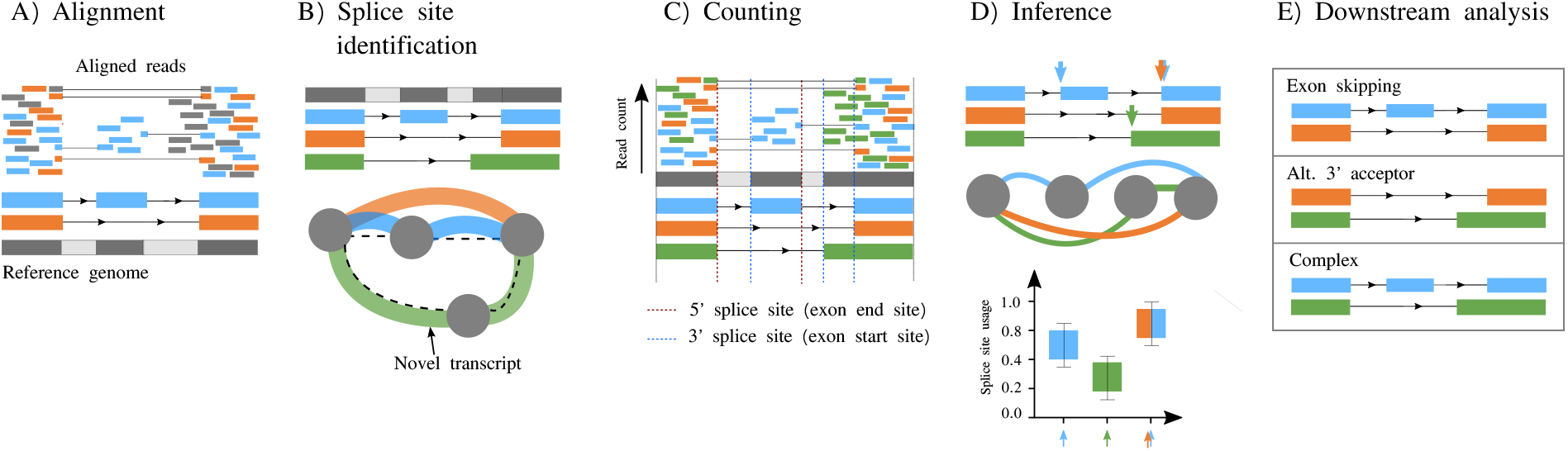
McSplicer workflow summary. The main steps of the McSplicer analysis are: A) Map RNA-seq reads to the reference genome sequence [10]. B) Identify annotated as well as novel splice sites through the reference-based assembly of transcripts using, e.g., StringTie [37]. C) Divide the gene into non-overlapping segments bounded by splice sites and count the number of reads mapping to distinct combinations of segments. D) Estimate splice site usages using McSplicer. E) Leverage splice site usages in various kinds of downstream analyses, such as the quantification of different types of alternative splicing events.

In the following sections, we introduce McSplicer’s model and algorithm for the estimation of parameters in that model. A more detailed description of the model and algorithms is provided in Supplementary Section 2. In the technical description of our model, we refer to the exon boundaries at the 3′ (acceptor) and 5′ (donor) splice sites as exon start and exon end sites, respectively. The description of our model is based on single-end reads which we apply to paired-end reads in Section 3.3. In the next section, we recapitulate the commonly assumed generative model of RNA-seq that also underlies the McSplicer model. For the sake of simplicity, we introduce the model based on individual observed reads and explain how parameters can be estimated from (much fewer) signature counts at the end of Section 2.3.

### 2.1 A generative model for RNA-seq reads

Consider the RNA-seq reads that mapped to a given gene. Reads are derived from one end of each of *N* fragments and each read has length *L*. We assume that each fragment is independently generated from one of the possible transcripts allowed by our model (see next Section). In this section, we describe a generative model for the sequence of the *n*-th read *R*_*n*_. The probability of *R*_*n*_ can be written as

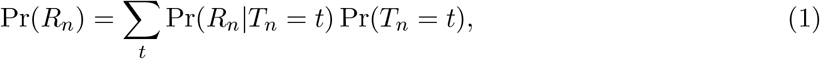

where *T*_*n*_ represents the transcript from which *R*_*n*_ was generated. Following models in [29] and [27], we assume that the probability of generating *R*_*n*_ from a transcript *t* is proportional to the product of the (effective) length [39] of the transcript, *l*(*t*), and the relative abundance of the transcript, *w*(*t*):

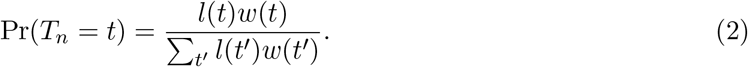

We introduce *B*_*n*_ that denotes the start position of *R*_*n*_ in *T*_*n*_, leading to

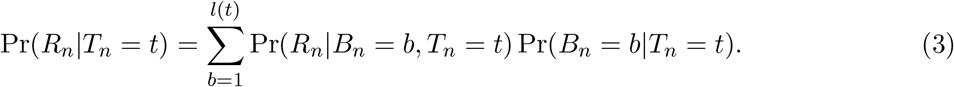

Assuming that *R*_*n*_ was generated uniformly across transcript *t*, we have

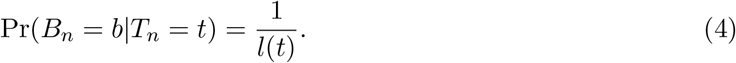

Pr(*R*_*n*_|*B*_*n*_ = *b*, *T*_*n*_ = *t*) = 1 if *R*_*n*_ is identical to the sequence of length *L* starting at a position *b* in transcript *t*, and this probability is 0 otherwise.

### 2.2 McSplicer: an inhomogeneous Markov chain to model the relative abundance of transcripts

We propose a new model for the relative abundance of transcripts expressed by a gene, denoted by *w*(*t*) in the previous section. Suppose we have obtained in step (B) in the McSplicer workflow (Fig. 2) potential exon start sites, 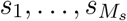, and potential exon end sites, 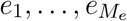, ordered by their occurrence in forward direction of a given gene. These splice sites partition the gene into non-overlapping segments *X*_1_, …, *X_M_*, where *M* = *M*_*s*_ + *M*_*e*_ + 1 and each segment is defined by a region between a pair of consecutive splice sites with the exception of the first and last segments (see Fig. 2C and Fig. 3). We introduce a sequence of hidden variables, *Z* = (*Z*_1_, …, *Z*_*M*_), where *Z*_*i*_ is a binary indicator for whether the *i*-th segment *X_i_* is transcribed (*Z_i_* = 1). Then, a particular transcript can be represented by a sequence of states for *Z*, as illustrated for transcripts *t*_1_, *t*_2_, *t*_3_ in Fig. 3. Thus, we can model the relative abundance of transcripts by modelling the probability of *Z*.

**Figure 3:**
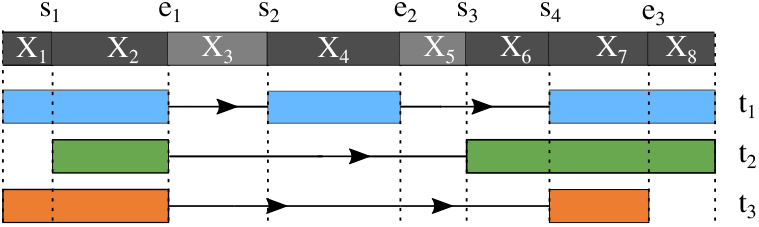
Example of hidden states representing 3 different transcripts. The gene includes four potential exon start sites and three potential exon end sites. These splice sites divide the gene into eight segments (*M*_*s*_ = 4, *M*_*e*_ = 3, and *M* = 8). The three sequences of states of *Z*, (1, 1, 0, 1, 0, 0, 1, 1), (0, 1, 0, 0, 0, 1, 1, 1), and (1, 1, 0, 0, 0, 0, 1, 0), represent the three transcripts *t*_1_, *t*_2_, and *t*_3_, respectively.

We use an inhomogeneous Markov chain to model the probability of the sequence of hidden variables, *Z* = (*Z*_1_, …, *Z*_*M*_). Specifically, the initial probability is given by

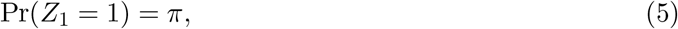

where *π* represents the proportion of transcripts that contain the first segment. We model the transition probability from *Z*_*i*_ to *Z*_*i*+1_ for *i* = 1, …, *M* − 1 as follows. If two consecutive segments *X*_*i*_ and *X*_*i*+1_ are separated by an exon start site *s*_*m*_,

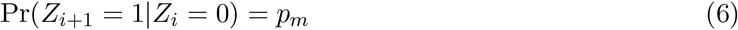

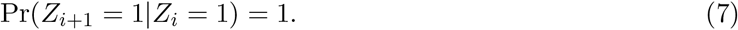

If they are separated by an exon end site *e*_*m*_,

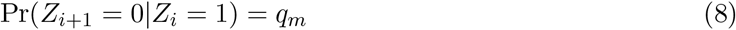

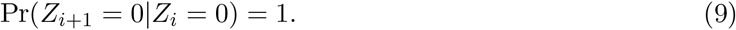

That is, if the current segment is transcribed (*Z*_*i*_ = 1), the splicing process ignores an exon start site (Equation 7), but it considers the potential usage of an exon end site *e*_*m*_ and decides to use it, i.e. end the exon, with its usage probability *q_m_* (Equation 8). On the other hand, if the current segment is not transcribed (*Z*_*i*_ = 0), the splicing process ignores an exon end site (Equation 9), but it uses an exon start site *s*_*m*_ with its usage probability *p*_*m*_ (Equation 6). The parameters 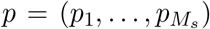 and 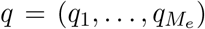 represent probabilities of using the corresponding splice sites given that each site is considered for potential usage. Throughout the rest of this work, we refer to these usage probabilities simply as usages. Table 1 shows the relative abundances defined by the proposed model for the three transcripts presented in Fig. 3. A more detailed description is provided in Supplementary Sections 2.1 to 2.3.

**Table 1:**
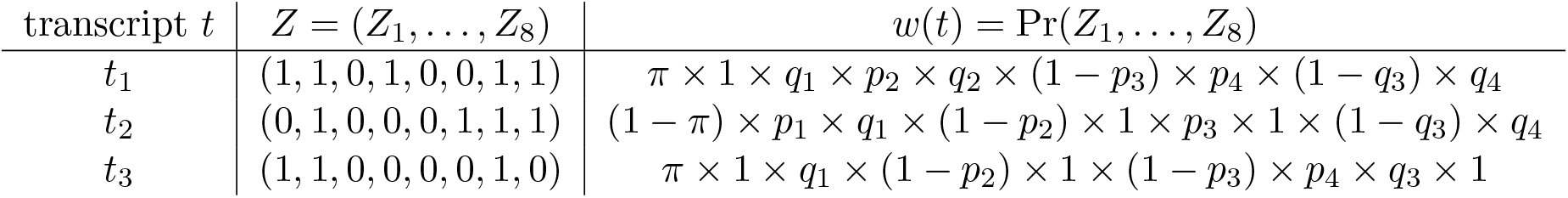
The relative abundances defined by the McSplicer model for the three transcripts presented in Fig. 3.

### 2.3 Parameter estimation and uncertainty quantification

We use an EM algorithm to compute the maximum likelihood estimates for the model parameters Θ = {*π, p, q*}, that is 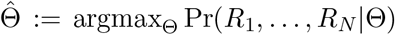. The complete log likelihood in the EM algorithm is 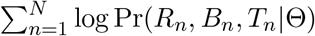. By combining the generative model and the McSplicer model in the previous two sections, Pr(*R*_*n*_, *B*_*n*_ = *b*, *T*_*n*_ = *Z*|Θ) for *b* ∈ {1, …, *l*(*Z*)} can be written

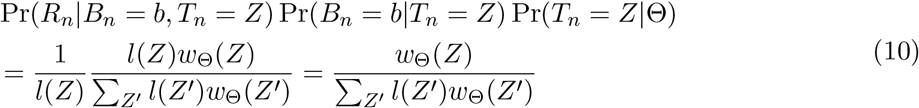

if *R*_*n*_ is identical to the sequence of length *L* starting at position *b* in transcript *Z*. Otherwise, this probability is 0. The details of the application of the EM algorithm to the proposed model are provided in Supplementary Section 2.4.2. The EM algorithm uses several quantities that we compute using dynamic programming, see Supplementary Section 2.4.1. Also, all quantities required in our EM algorithm can be computed only using signature counts (Supplementary Section 2.4.2), so the input to the McSplicer are the signature counts rather than individual reads.

We quantify the uncertainty of our estimator 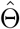 using bootstrapping. Specifically, let 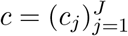 represent the signature counts over *J* signatures defined for a given gene, where the total signature count equals the total read count in the gene, i.e., 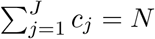. We draw *B* independent bootstrap samples, *c*^1^, …, *c*^*B*^, from a multinomial distribution:

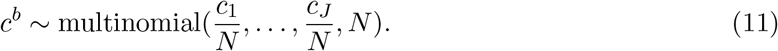

Then, we compute *B* bootstrap estimators, 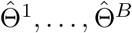, by applying our EM algorithm to each bootstrap sample and use them to approximate the sampling distribution of our estimator 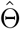. In this paper, we quantify the uncertainty of 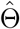 using a confidence interval computed from the approximated sampling distribution. Other types of uncertainty quantification could easily be obtained from the bootstrap estimators.

## 3 Results

We assess the performance of McSplicer in comparison to existing state-of-the-art methods on both simulated and real RNA-seq data sets. Simulated data allow to compare estimates to a known ground truth of expressed transcripts and thus known quantities of alternative splicing events. On the other hand, simulated data cannot fully capture the complexity of data sets generated in real RNA-seq experiments.

We compare the performance of McSplicer to SplAdder, MAJIQ, and StringTie. In Supplementary Section 2.5 we provide details on software versions and command line arguments used. SplAdder was used in a large-scale study [22] to detect and quantify alternative splicing events in nearly 9,000 tumor RNA-seq samples. In a comparative benchmark analysis performed in [23], it showed a better performance than competing methods JuncBase [6], rMATS [42], and SpliceGrapher [40]. Compared to SplAdder, which is limited to the detection of simple types of splicing events, MAJIQ introduced a novel approach that additionally captures more complex transcript variations. MAJIQ was shown in a recent benchmark [31] to compare favorably to existing state-of-the-art methods and the authors demonstrated in [49] that MAJIQ also outperforms LeafCutter and rMATS.

StringTie, on the other hand, assembles and quantifies full-length transcripts from RNA-seq but was not specifically designed for the quantification of splice site usage. Nevertheless, splice site usage can be inferred from the abundance of the assembled transcripts and we include this approach as a baseline in our benchmark: In all experiments, McSplicer uses StringTie to construct the exonintron structure in steps (B) and (C) of the workflow (Fig. 2), which potentially contains novel splice sites. In contrast to the inference of splice site usage from expressed full-length transcripts, however, McSplicer estimates the usage of the same set of splice sites using the EM algorithm described in the previous section.

### 3.1 Comparable splice sites

Each method, however, uses a different set of parameters to quantify alternative splicing events. SplAdder quantifies four canonical types of splicing events using the widely used *percent spliced in* (PSI) metric. PSI denotes the ratio between the number of reads supporting one outcome of the event (e.g the inclusion of an exon) over the number of reads directly supporting either of the two alternative outcomes. Similarly, MAJIQ computes the *percent selected index* (Ψ) for each splice junction involved in a *local splicing variation* (LSV), which denotes its fractional usage. To ensure a meaningful comparison of splice site usages in McSplicer to PSI from SplAdder and Ψ from MAJIQ, we only consider splice sites for which the meaning of these three quantities, if defined, coincide. These *comparable* splice sites are obtained from alternative splicing events between two transcripts such that all remaining transcripts expressed by a gene consistently support one of the two possible outcomes of the event.

More formally, let *s*_1_, *s*_2_, …, *s*_*M*_ denote the splice sites of a gene *G*, ordered by their genomic coordinates. Consistent with [14], we define *alternative splicing events* for pairs of transcripts *t*_1_, *t*_2_ as maximal sequences *s*_*i*_, …, *s*_*j*_ of alternative splice sites, i.e. splice sites that are used by *t*_1_ or by *t*_2_, but not by both. To distinguish the outcome of alternative splicing from the outcome of alternative transcription initiation or termination, we additionally require that *s*_*i*−1_ and *s*_*j*+1_ denote common donor and acceptor sites, respectively. If every transcript expressed by *G* is consistent with *t*_1_ or *t*_2_ in its use of *s*_*i*−1_, *s*_*i*_, …, *s*_*j*+1_, we call the alternative splice sites *s*_*i*_, …, *s*_*j*_ *comparable*. Note that the definition of comparable splice sites is invariant with respect to the choice of *t*_1_ and *t*_2_ among expressed transcripts of *G*. For an illustrative example of comparable and non-comparable splice sites see Supplementary Figure S3.

For comparable splice sites of simple events, the three different parameters, i.e. splice site usage, PSI and Ψ, equally reflect the relative abundance of transcripts expressed by a given gene that use the splice site, or equivalently contain the corresponding exon. Analogously, Ψ and splice site usage are equivalent for comparable splice sites of complex events. From StringTie assemblies of full-length transcripts, estimates of splice site usage can directly be obtained from the relative abundance of transcripts using a given splice site.

### 3.2 Simulation study

We used Polyester [15] to simulate reads from a human transcriptome with abundances estimated from a real RNA-experiment (GEO accession GSM3094221) using RSEM [28]. Based on these ground truth expressions, we simulated data sets with varying sequencing depth commonly observed in practice, including 20 million, 50 million, and 75 million reads of 100bp length. Following the same strategy as [44], we randomly selected a set of 1000 genes with at least two expressed transcripts and sufficiently high ground truth expression (gene-level read count per kilobase above 500). Among comparable splice sites we exclude from the analysis constitutive ones with usage 1 and splice sites that are not used by any of the expressed transcripts (usage 0).

From the ground true abundance of transcripts, we calculate for each comparable splice site its true usage as the relative contribution of transcripts using a given splice site to the total expression of a gene. Specifically, let *A*(*s*) and *B*(*s*) denote subsets of transcripts in a gene *G* that either use or do not use a particular splice site *s*, respectively. Then the true usage of splice site *s* is computed by

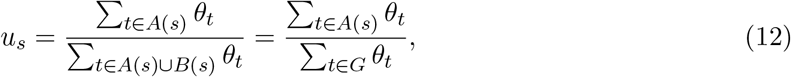

where *θ*_*t*_ represents the true abundance of transcript *t*. Since for comparable splice sites the true usage is equivalent to the true PSI value or the true Ψ, we will consistently refer to them in the following as splice site usage.

We quantify the accuracy of splice site usages inferred by each method by using the Kullback-Leibler (KL) divergence. For a given splice site *s*, the two possible outcomes, whether or not a transcript uses the splice site can be modelled by a Bernoulli distribution with the splice site usage *u*_*s*_, denoted by Bern(*u*_*s*_). Let 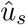 represent the estimated splice usage. Then, we measure the accuracy of 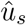 using the KL divergence of Bern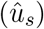 from Bern(*u*_*s*_):

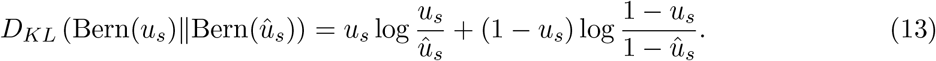

#### 3.2.1 McSplicer more accurately infers splice site usage than competing methods

We distinguish splice sites by the type of event they are part of, including exon skipping, intron retention, alternative 3′ and 5′ splice sites, and complex events that cannot be assigned to one of the canonical types. The events are labeled by Astalavista [14] through a pairwise structural comparison of all transcript species expressed in our ground truth transcriptome (see Supplementary Figures S1 and S4).

The number of variable (i.e. usage 0 < *u* < 1) comparable splice sites in our simulated data set with corresponding event types defined by Astalavista is listed in Supplementary Table S1. Supplementary Table S1 also lists the total number of splice sites per type reported by all four methods. While McSplicer will quantify the usage of all splice sites except those missed by StringTie in step (B) in Fig. 2, competing methods report only events that satisfy an adjustable confidence threshold (SplAdder) or are considered reliable according to internal filters (MAJIQ). As a result, both MAJIQ’s and SplAdder’s accuracy is evaluated on a smaller, presumably more confidently estimated set of events (Supplementary Table S1) and are otherwise not penalized for missing events.

MAJIQ estimates two parameters that correspond to the relative usage of a skipped exon, one based on the intron connecting it to the upstream exon, and one based on the downstream exon (Supplementary Figure S5). Here, we compare the performance to the latter one, which we observed to be slightly more accurate. The former is reported in Supplementary Figure S6.

Fig. 4 compares the accuracy of splice site usages inferred by McSplicer and competing methods from 50 million reads on four canonical types of events as well as on complex events. For each method, only events reported and quantified by that methods are considered. Supplementary Figure S7 shows consistent results when considering events that McSplicer and competing methods have pairwise in common. Across all types of events, McSplicer infers splice site usages more accurately than competing methods. The accuracy of splice site usage inferred by McSplicer is not affected by the complexity of the event, whereas MAJIQ’s estimates are substantially less accurate for complex events. SplAdder is restricted to the quantification of simple events. As originally reported by the authors in [23], SplAdder quantifies intron retentions less accurately than other simple types of events. Other methods, including McSplicer, perform well on this type of event, which plays an important role for cell development in mammals [4], is commonly observed in cancerous tissues [12], and is a source of neoepitopes in cancer [43]. Compared to baseline splice site usage extracted from StringTie transcript assemblies, McSplicer utilizes StringTie’s transcript models to substantially refine the quantification of local splicing variation. Similar results were obtained on data sets comprising 20 million and 75 million reads (see Supplementary Figure S8 and Figure S9).

**Figure 4:**
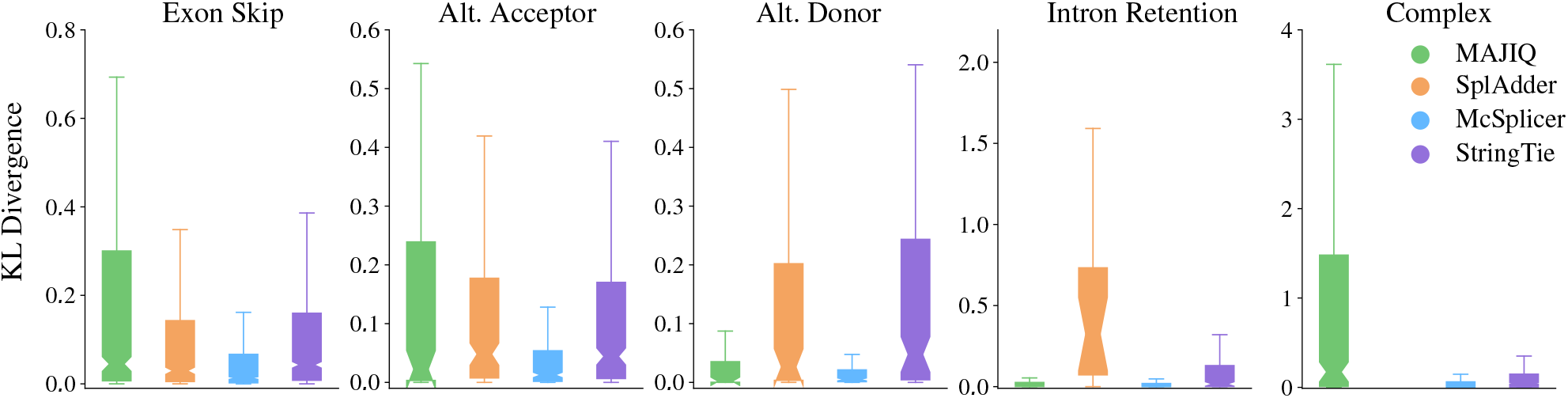
Accuracy of McSplicer and competing methods in quantifying the usage of variable splice sites from 50 million simulated RNA-seq reads. For each method, only splice sites in events that the method reports and quantifies are considered. SplAdder is limited to the quantification of simple AS events.

#### 3.2.2 McSplicer leverages all reads mapped to a gene

McSplicer makes use of all reads mapped to a given gene to simultaneously infer parameters in the McSplicer model, while other methods typically use only reads that directly support their parameters.

To quantify the contribution of the simultaneous inference in McSplicer to improve the accuracy of estimators, we estimate one splice site usage parameter at a time using only reads directly supporting the parameter. Similar to the calculation of the traditional PSI metric, we remove for each event with comparable splice sites all reads that do not overlap any of the event’s exons, and run and evaluate McSplicer on the resulting restricted instance as described in the previous section. Fig. 5 confirms that McSplicer profits enormously from transcriptional evidence that lies outside of the local splicing event. Across all types of events, McSplicer estimates splice site usage less accurately when reads that do not overlap an event are removed.

**Figure 5:**
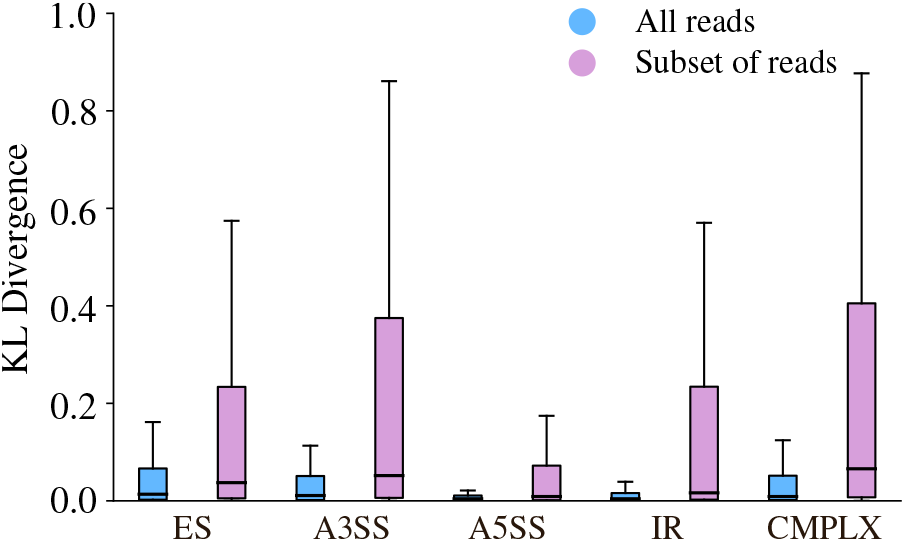
McSplicer leverages all RNA-seq reads mapped to a gene to improve the accuracy of splice site usage estimates. McSplicer achieves lower KL divergence form true splice site usages when considering all reads mapped to a gene locus at once (blue) compared to using only reads that overlap any of the event’s exons (pink). ES denotes exon skipping, A3SS alternative 3’ splice site, A5SS alternative 5’ splice site, IR intron retention, and CMPLX complex events.

### 3.3 McSplicer estimates agree with Spike-In RNA Variants

To evaluate the performance of McSplicer under the full complexity of data derived from a real RNA-seq experiment, we used spike-in controls that were previously added to human monocyte-derived macrophages [20] from five different donors (GEO accession number GSE117206). The Spike-In RNA Variants (SIRV) [36] comprise 69 synthetic RNA molecules that were added in known relative concentrations before library preparation. Mimicking the complexity of 7 human model genes, between 6 and 18 artificial transcripts per gene vary in different types of alternative splicing, transcription start- and end-sites, or are transcribed from overlapping genes, or the antisense strand. The concentration ratios between different SIRV isoforms span a range of more than two orders of magnitude. For each donor sample, including artificial SIRV isoforms, Hoss et al. [20] sequenced 200 million paired-end reads of 2 × 125bp length. McSplicer considers both mates independently as inputs reads *R*_*n*_ (see Section 2.3).

Leveraging the artificial reference genome (SIRVome) and the known relative mixing ratios of SIRV isoforms, we derive ground truth splice site usages (Equation 12). Again, we obtain event labels from Astalavista, which comprise 26 variable splice sites in simple events and 12 in complex events. In this experiment, we do not restrict the evaluation to comparable splice sites but include all variable sites. Fig. 6 compares splice site usages as estimated by McSplicer to the true usages in one of the five samples (donor 5). A Spearman’s rank correlation coefficient of *ρ* = 0.798 indicates a good agreement between estimated and true usages. We obtain similar results on the remaining four samples (Supplementary Figure S10).

**Figure 6:**
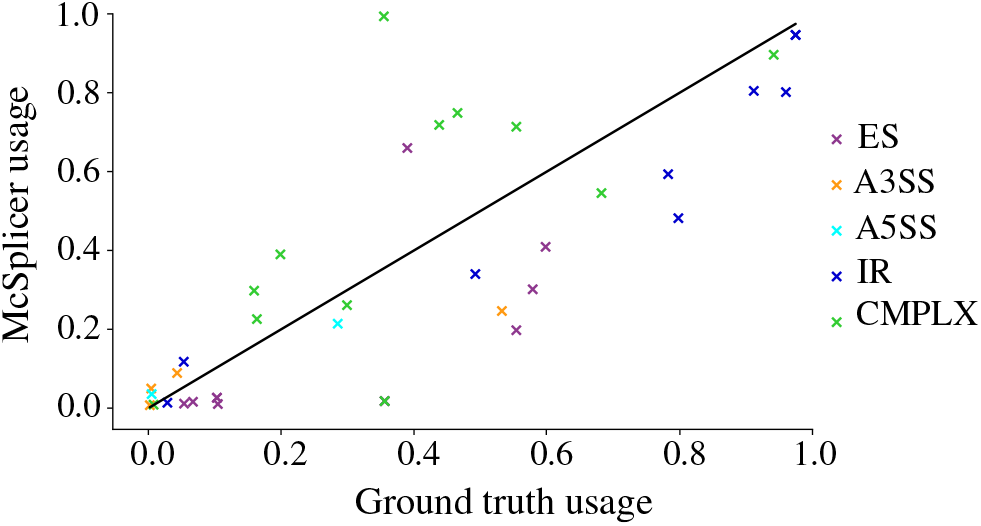
McSplicer results on spike-in RNA variants (SIRV), donor sample 5. Ground truth splice site usages computed from known mixing ratios of SIRV isoforms are compared to usages estimated by McSplicer. Out of 38 variable splice sites, 26 belong to simple events and 12 belong to complex events. ES denotes exon skipping, A3SS alternative 3’ splice site, A5SS alternative 5’ splice site, IR intron retention, and CMPLX complex events.

SplAdder and MAJIQ only report between 6 and 12 among all 38 true events, too few to allow for a meaningful quantification of agreement between estimated and true PSI and *ψ* values. Supplementary Figures S11 and S12 show the corresponding scatter plots for PSI and *ψ* values estimated by SplAdder and MAJIQ, respectively.

### 3.4 Quantifying the effect of cryptic splice site mutations in patients with autism spectrum disorder

In this section, we illustrate the utility of splice site usages computed by McSplicer in interpreting the potentially complex effect of genetic variants on RNA splicing. Non-coding genetic variants that alter mRNA splicing play an important role in rare genetic diseases [8]. In [21], the authors use a deep neural network to identify non-coding genetic variants that disrupt mRNA splicing. They identified a set of high-confidence *de novo* mutations predicted to disrupt splicing in individuals with intellectual disability and individuals with autism spectrum disorders (ASD). To validate them, the study included RNA-seq experiments (270-388 million 150bp reads per sample) of peripheral blood-derived lymphoblastoid cell lines from 36 individuals with ASD. Based on the presence of reads spanning the corresponding splice junction, the authors validate 21 aberrant splicing events associated with the predicted *de novo* mutations. Each of the splicing events was uniquely observed in one individual.

Here, we employ McSplicer to quantify the effect size of the mutations associated with the aberrant splicing events in ASD patients based on RNA-seq data. In fact, in a different experiment in [21], the authors computed the effect size by comparing the relative usage of novel splice sites between individuals with and without the mutations. Relative usage of a novel donor site was calculated based on the number of reads supporting the junction to the closest annotated acceptor. The annotated donor closest to the novel one was selected as reference splice site. Relative usage of novel acceptors were calculated analogously. In [21] the authors point out, however, that computing the effects size of splicing mutations based on a pre-selected set of incident splice junctions likely underestimates the true effect size since, among other shortcomings, not all isoform changes are taken into account.

In contrast, McSplicer’s model of splice site usage does not depend on an ad hoc selection of specific junctions or AS units but naturally captures simultaneous changes in expression of multiple isoforms expressed by a gene. We therefore utilized McSplicer to quantify the effect size of the validated *de novo* mutations on splice sites in ASD patients. We excluded 11 aberrant splicing events where only 1 or 2 spliced reads supported the novel splice site or junction. For each *de novo* mutation and the corresponding aberrant splicing event, we used McSplicer to estimate splice site usage and to compute 95% bootstrapping confidence intervals for the individual harboring the variant and a control individual with similar sequencing depth. For all 10 aberrant splicing events, we observe significantly different splice site usages (i.e., the two confidence intervals do not overlap) between mutated and control ASD individuals (Supplementary Table S2). Fig. 7 provides three illustrative examples. For gene *ENOPH1*, McSplicer estimates a decrease in usage of the acceptor site directly affected by the variant, consistent with the increased skipping of the corresponding exon that can be observed in the Sashimi plot. In gene *CORO1B*, a novel donor site is used exclusively in the individual with the variant, identified and quantified with non-zero usage by McSplicer. For gene *PCSK7*, McSplicer estimates a decrease in usage of the affected donor sites, consistent with the retention of the downstream intron.

**Figure 7:**
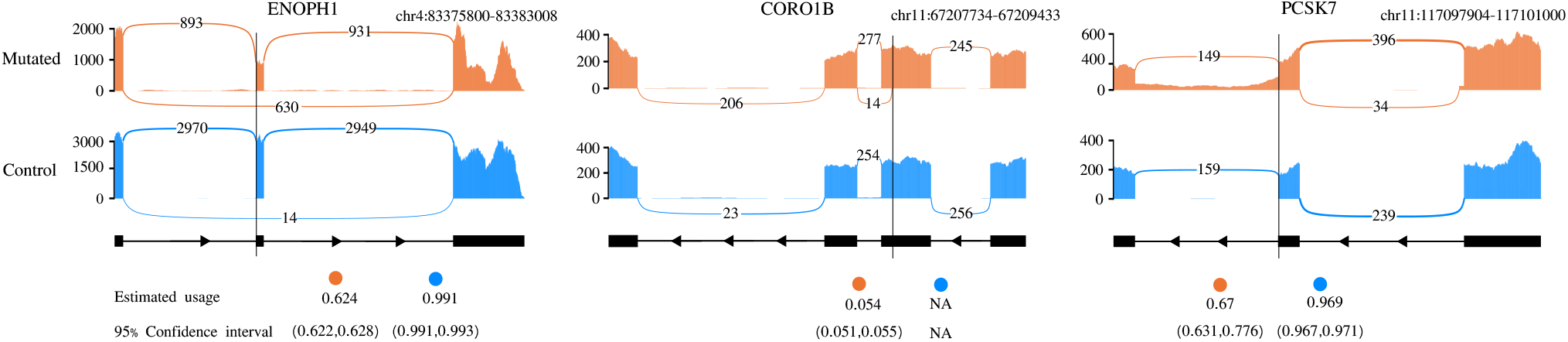
McSplicer splice site usage estimates and 95% bootstrapping confidence intervals for three disrupted splicing events reported in ASD patients versus control individuals. Variant locations are indicated by black vertical lines. Each plot illustrates the gene structure around the event with the precise genomic window specified on top, the read coverage and the junction read count. The Sashimi plots shown here are created using the ggsashimi tool [16].

## 4 Conclusion

We have introduced McSplicer, a novel method that estimates splice site usage across expressed transcripts. Rather than attempting to reconstruct expressed transcripts, McSplicer is based on a simplified probabilistic splicing model that has generated the set of expressed transcripts. It is not restricted to a pre-defined class of alternative splicing events or units but our probabilistic model is able to describe arbitrarily complex types of splicing patterns based on few, easy to interpret, parameters. We estimate these parameters, i.e. splice site usages, using all read data at once and demonstrate in experiments that this yields more accurate estimates compared to other methods that use only reads directly supporting their parameters. Through its integration with transcript assembly methods such as StringTie, McSplicer quantifies the usage of annotated as well as novel splice sites.

Our model for relative transcript abundance assumes the Markovian property across indicators (*Z*_*i*_) for whether a segment is transcribed. This assumption allows for an efficient algorithm to estimate parameters of the model, but it potentially limits the ability of our method to model longer range dependencies such as between the recognition of 5’ and 3’ splice sites or between the removal of introns within transcripts. If true dependencies are longer than our model can describe, the individual estimators for splice site usages may still be accurate, but we expect transcript frequencies implied by our model to be less accurate [27]. One way to model longer range dependencies is to use higher order Markov chains as long as the data provide sufficient information to estimate these dependencies.

The splice site usages computed by McSplicer can be leveraged in various types of downstream analyses, such as the statistical comparison of splice site usage between different conditions [30], the quantification of various types of splicing events, the identification of subgroups of samples that show similar splicing patterns (i.e., unsupervised clustering [32]), or the discrimination between alternatively spliced and constitutive exons [35].

We have used McSplicer to quantify the effect size of splicing mutations in ASD patients. In this context, splice site usage as computed by McSplicer can be considered analogous to the “strength” of a splice site predicted by methods such as SplicePort [11] from sequence-based features. Point mutations in the consensus splice site sequence can affect the strength of a splice site and result in the skipping of the exon or even the skipping of multiple exons [47], or the activation of cryptic splice sites in monogenic disorders [3]. In fact, a single nucleotide substitution might produce multiple (erroneous) splicing isoforms at the same time, as has been observed, for example, for specific mutations in patients with cystic fibrosis (3 isoforms) [38], Ehlers-Danlos syndrome (4 isoforms) [47], Duchenne muscular dystrophy (3 isoforms) [17], and X-linked spondyloepiphyseal dysplasia tarda (7 isoforms) [53]. McSplicer does not attempt to reconstruct every single aberrant isoform, but similar to a weakening (strengthening) of a splice site as predicted from sequence alterations by, e.g., the Shapiro splice site probability score [41], the effect of a mutation will be reflected in a reduced or increased usage of the corresponding splice site estimated from RNA-seq reads.

In our analysis of ASD patients data, we computed the effect size for each mutation associated with an aberrant splicing event by separately estimating splice site usages in two individuals with and without the variant, and then computing the difference between the estimated usages. We quantified the significance of the estimated effect size by checking whether the confidence intervals for the two estimated usages overlap. This procedure does not use the full data from multiple individuals and fails to consider variability among individuals, possibly leading to an increased number of false positives. Methods that model differences in splice site usages between individuals from multiple groups and exploit the variability among them should perform better in estimating effect size and quantifying their uncertainty.

## 5 Acknowledgments

We thank Matthew Stephens for invaluable discussions on building the proposed model and Zhen Zuo for help with testing the method on other data sets. We thank the members of H. Shim, S. Canzar, and T. Speed groups for helpful comments. This work was supported by the Purdue startup fund. I.A. was supported by a Deutsche Forschungsgemeinschaft fellowship through the Graduate School of Quantitative Biosciences Munich.

