## Supplementary Material for "McSplicer: a probabilistic model for estimating splice site usage from RNA-seq data"

#### 1 Supplementary Figures and Tables

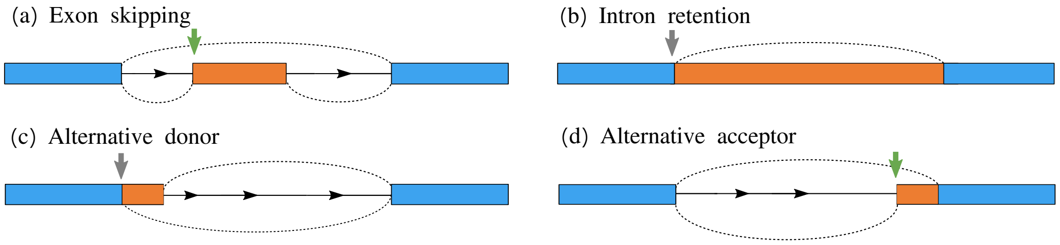


Figure S1: Four types of simple alternative splicing events. Blue rectangles represent constitutive exon or exonic segments. Orange rectangles represent alternatively spliced ones. (a) The usage of the marked acceptor site defines the relative abundance of the inclusion of the skipped exon. (b) For intron retentions, the usage of the marked splice site defines the relative abundance of the inclusion of the intron. (c) For alternative donors (c) and alternative acceptors (d), the usage of the marked donor and acceptor sites determine the relative abundance of the two alternative events.

*
†

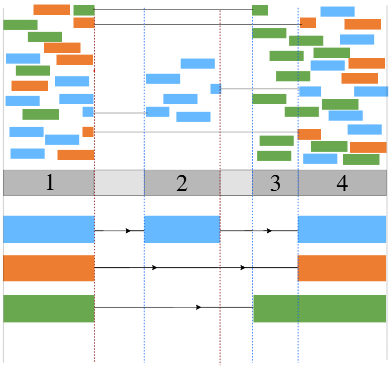


| signature | signature count |
|---|---|
| {1} | 16 |
| {1 − 2} | 1 |
| {1 − 3} | 1 |
| {1 − 4} | 2 |
| {1 − 2 − 4} | 0 |
| {1 − 3 − 4} | 0 |
| {2} | 4 |
| {2 − 4} | 1 |
| {3} | 4 |
| {3 − 4} | 4 |
| {4} | 15 |

Figure S2: An illustrative example showing signatures with their corresponding read counts. McSplicer estimates splice site usage from these *signagture counts* rather than from individual read alignments. In this example, three transcripts imply a partitioning into 4 non-overlapping exonic segments. Read colors indicate the originating transcript.

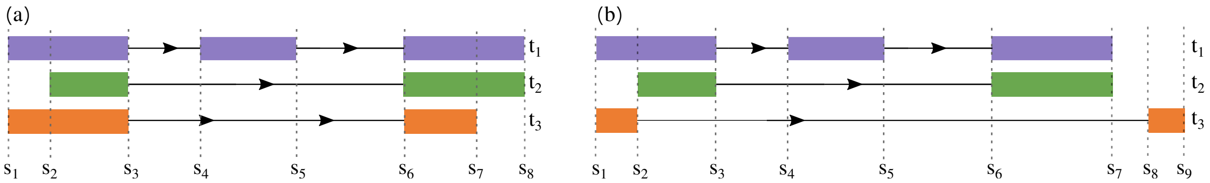


Figure S3: (a) An example of an exon skipping event involving *comparable* splice sites. The splice sites $s_4$ and $s_5$ are used exclusively by $t_1$, but splice sites $s_3$ and $s_6$ are common donor and acceptor sites to all three transcripts $t_1$, $t_2$, and $t_3$. (b) An example of an exon skipping event with non-comparable splice sites. The splice sites $s_4$ and $s_5$ are used exclusively by $t_1$. The splice sites $s_3$ and $s_6$ denote the common donor and acceptor sites of $t_1$ and $t_2$, Transcript $t_3$, however, is inconsistent with both $t_1$ and $t_2$ in its use of splice sites $s_3 \ldots s_6$.

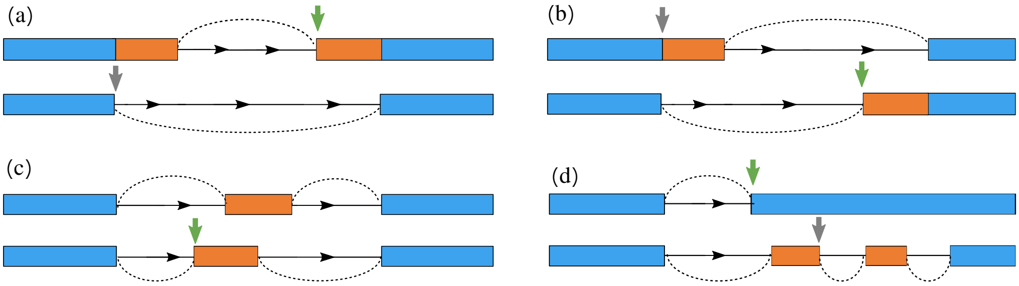


Figure S4: Examples of complex patterns of alternative splicing [2]. Green and grey arrows highlight the corresponding varying $3'$ and $5'$ splice sites, respectively. These are illustrative examples of the varying non-redundant splice sites which we consider in our benchmark.

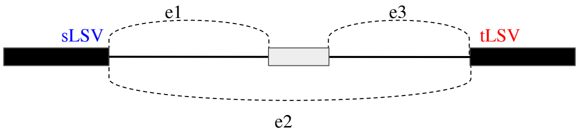


Figure S5: MAJIQ computes the percent selected index ($\psi$) for each junction involved in a local splicing variation (LSV) which denotes its fractional usage. An exon skipping event can be inferred either from the estimated $\psi$ value of edge $e_1$ connecting source LSV (*sLSV*) to the cassette exon, or edge $e_3$ connecting the cassette exon to the target LSV (*tLSV*). We notice that the estimated usage $E[PSI(e_3)]$ tends to be slightly more accurate than $E[PSI(e_1)]$.

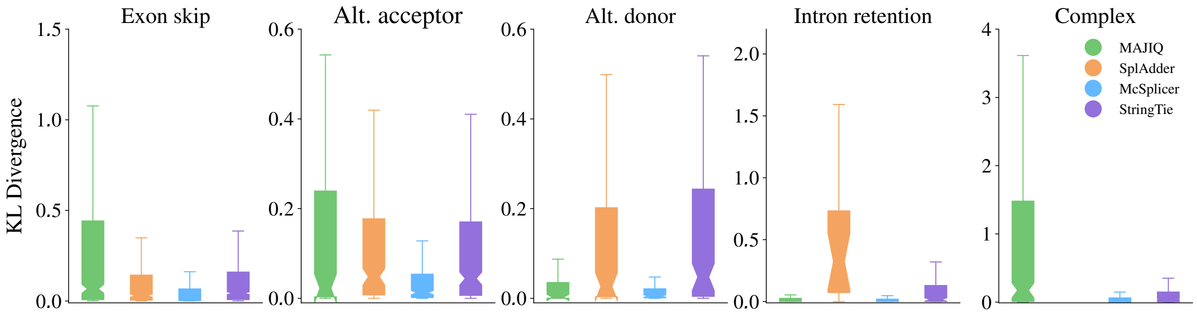


Figure S6: Accuracy of McSplicer and competing methods in quantifying the usage of variable splice sites from 50 million simulated RNA-seq reads. For MAJIQ, here we consider the estimated $\psi$ value of the edge incident to the source LSV. See Fig. S5 for an illustration.

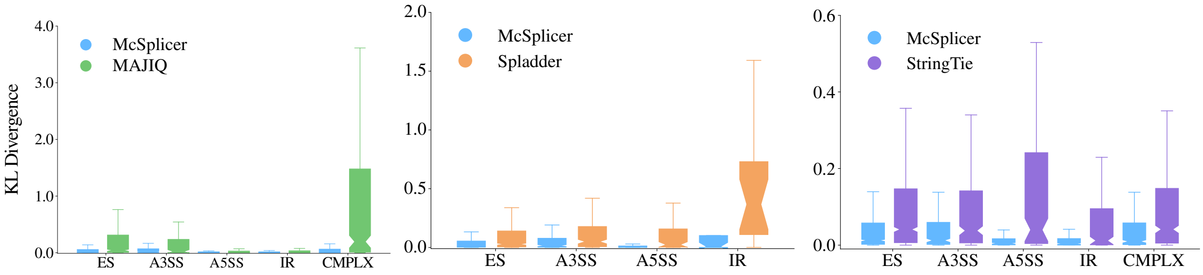


Figure S7: Accuracy of McSplicer and competing methods in quantifying the usage of variable splice sites from 50 million simulated RNA-seq reads. Events that McSplicer and competing methods have pairwise in common are considered. SplAdder is limited to the quantification of simple AS events.

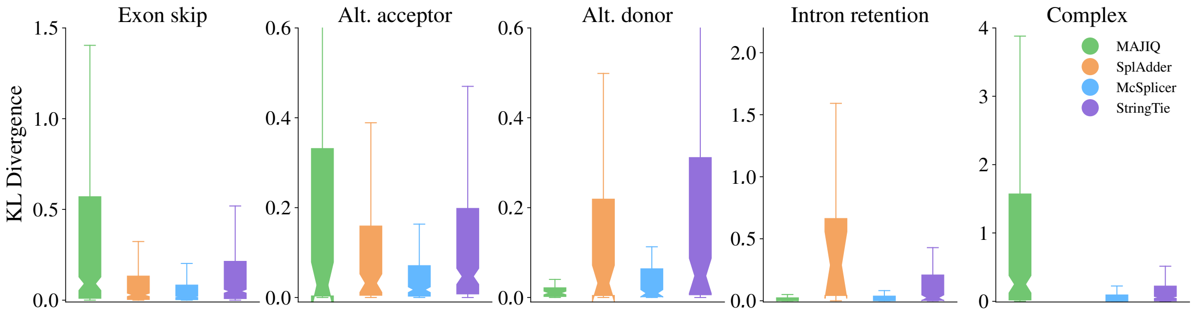


Figure S8: Accuracy of McSplicer and competing methods in quantifying the usage of variable splice sites from 20 million simulated RNA-seq reads. For each method, only splice sites in events that the method reports and quantifies are considered. SplAdder is limited to the quantification of simple AS events.

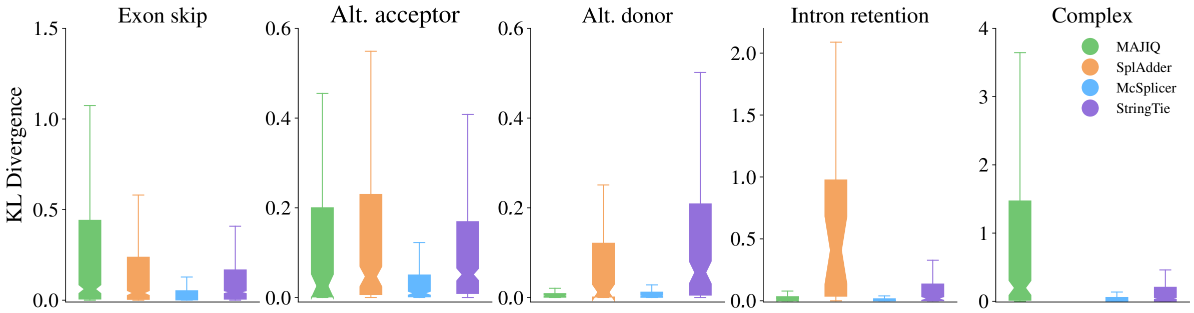


Figure S9: Accuracy of McSplicer and competing methods in quantifying the usage of variable splice sites from 75 million simulated RNA-seq reads. For each method, only splice sites in events that the method reports and quantifies are considered. SplAdder is limited to the quantification of simple AS events.

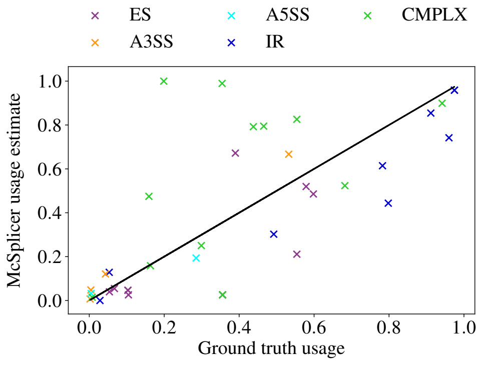


(a) Donor 1.
Spearman's $\rho = 0.774$.

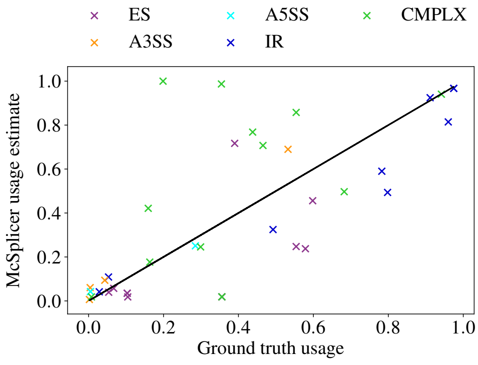


(b) Donor 2.
Spearman's $\rho = 0.769$.

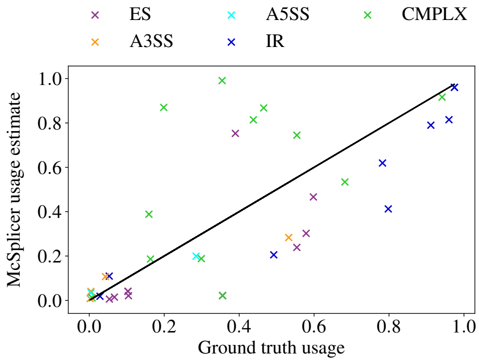


(c) Donor 3.
Spearman's $\rho = 0.774$.

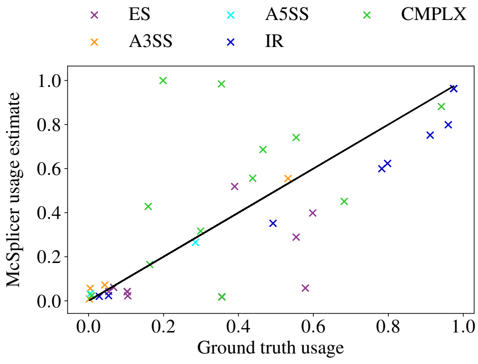


(d) Donor 4.
Spearman's $\rho = 0.782$.

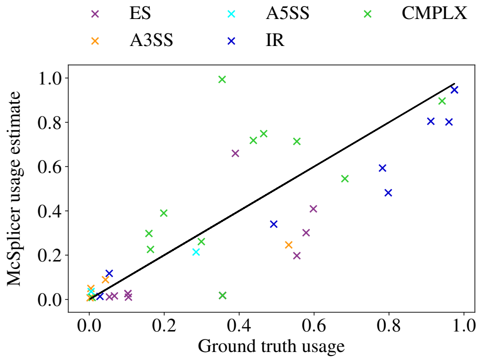


(e) Donor 5.
Spearman's $\rho = 0.798$.

Figure S10: McSplicer results on spike-in RNA variants (SIRV) on 5 different SIRV samples. Ground truth splice site usages computed from known mixing ratios of SIRV isoforms are compared to usages estimated by McSplicer. Out of 38 variable splice sites , 26 belong to simple events and 12 belong to complex events. ES: exon skipping; A5SS: alternative 5' splice site; A3SS: alternative 3' splice site; CMPLX: complex event; IR: intron retention.

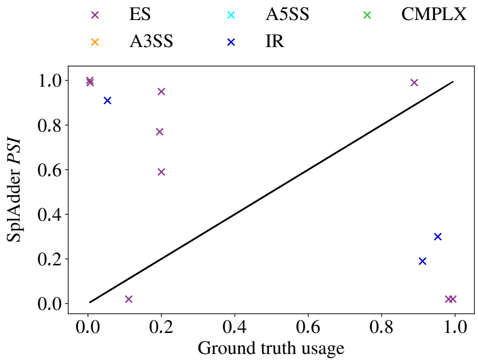


(a) Donor 1.

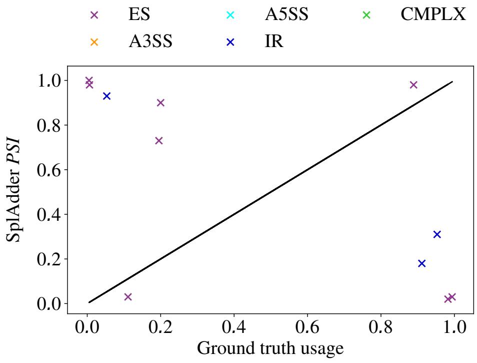


(b) Donor 2.

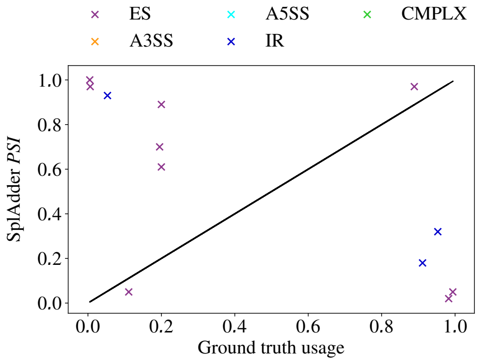


(c) Donor 3.

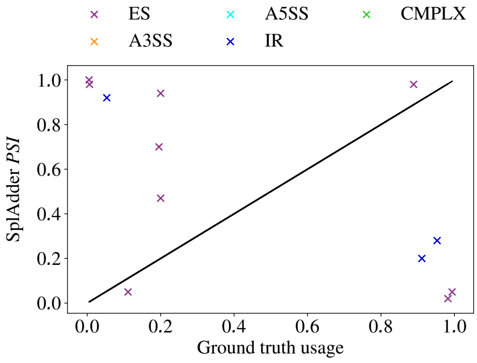


(d) Donor 4.

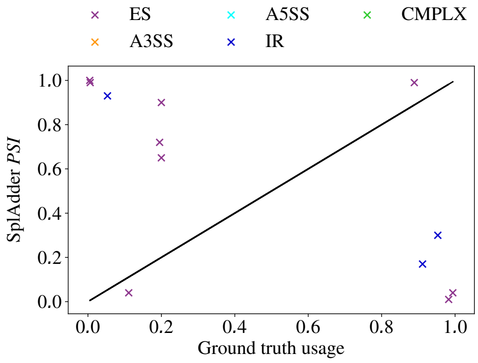


(e) Donor 5.

Figure S11: SplAdder results on spike-in RNA variants (SIRV) on 5 different SIRV samples. Ground truth splice site usages computed from known mixing ratios of SIRV isoforms are compared to usages estimated by SplAdder. Out of 38 variable splice sites , 26 belong to simple events and 12 belong to complex events. ES: exon skipping; A5SS: alternative 5' splice site; A3SS: alternative 3' splice site; CMPLX: complex event; IR: intron retention.

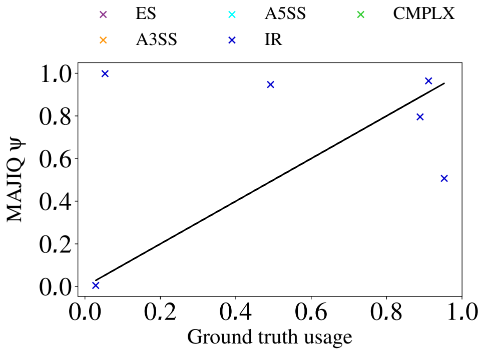


(a) Donor 1.

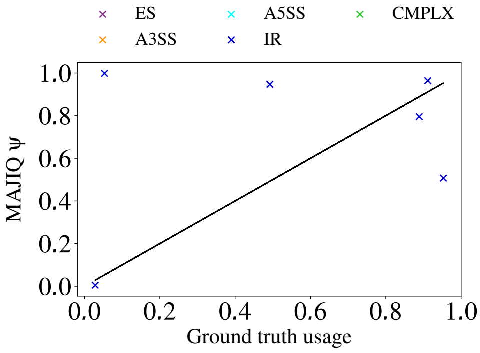


(b) Donor 2.

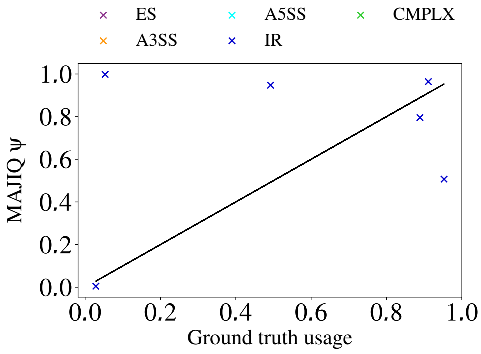


(c) Donor 3.

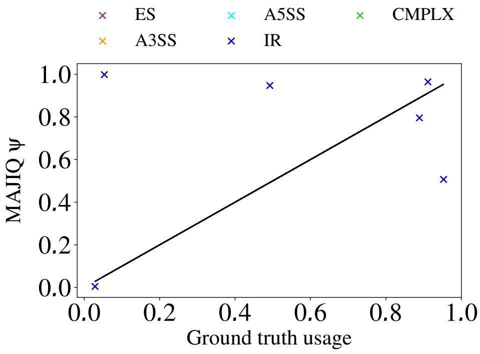


(d) Donor 4.

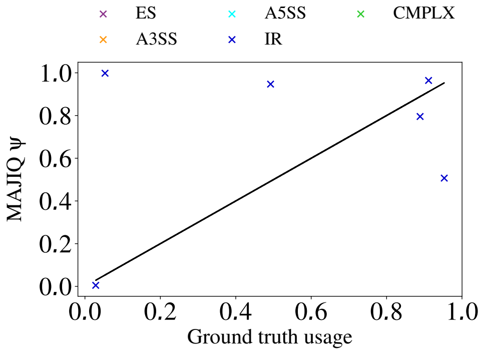


(e) Donor 5.

Figure S12: MAJIQ results on spike-in RNA variants (SIRV) on 5 different SIRV samples. Ground truth splice site usages computed from known mixing ratios of SIRV isoforms are compared to usages estimated by MAJIQ. Out of 38 variable splice sites , 26 belong to simple events and 12 belong to complex events. ES: exon skipping; A5SS: alternative 5’ splice site; A3SS: alternative 3’ splice site; CMPLX: complex event; IR: intron retention.

| | Exon skipping | Alt. 3' acceptor | Alt. 5' donor | Intron retention | Complex |
|---|---|---|---|---|---|
| AStalavista | 475 | 229 | 129 | 134 | 508 |
| SplAdder | 366 | 150 | 87 | 25 | - |
| MAJIQ | 371 | 106 | 81 | 89 | 429 |
| StringTie | 455 | 209 | 120 | 127 | 487 |
| McSplicer | 455 | 209 | 120 | 127 | 487 |

Table S1: The first row shows the number of comparable splice sites in our 50M RNA-seq simulated dataset classified by type as labeled by AStalavista. Each simple event contains by definition one such splice site, while complex events involve one or two variable splice sites that are comparable. The following rows show the number of varying splice sites classified by event type as quantified by each of the four methods.

| Gene name | chr | Splice Site | Mutated | Control | Effect size | Event type |
|---|---|---|---|---|---|---|
| BCL7B | 7 | 72966572 | 0.786 (0.784,0.792) | 0.956 (0.953,0.960) | -1.06 | ES |
| ENOPH1 | 4 | 83378068 | 0.624 (0.622, 0.628) | 0.991 (0.991,0.993) | -0.20 | ES |
| YME1L1 | 10 | 27431414 | 0.353 (0.349,0.356) | 0.81 (0.810,0.813) | -0.36 | ES |
| PPP4R2 | 3 | 73112824 | 0.463 (0.459,0.466) | 0.951 (0.950,0.955) | -0.32 | ES |
| TMBIM6 | 12 | 50153004 | 0.887 (0.887,0.889) | 0.945 (0.948,0.949) | -0.05 | ES |
| IDUA | 4 | 997837 | 0.209 (0.208,0.211) | 0.0 | ∞ | Novel A5SS |
| CORO1B | 11 | 67208804 | 0.054 (0.051,0.055) | 0.0 | ∞ | Novel A5SS |
| SHPRH | 6 | 146266702 | 0.546 (0.529,0.582) | 0.0 | ∞ | Novel IR |
| PCSK7 | 11 | 117098932 | 0.67 (0.631,0.776) | 0.969 (0.967,0.971) | -0.16 | Novel IR |
| ELOVL1 | 1 | 43829994 | 0.200 (0.200,0.215) | 0.0 | ∞ | Novel IR |

Table S2: McSplicer splice site usage estimates on mutated and control Autism samples with 95% bootstrapping confidence intervals shown in parentheses. We compute the effect size using the difference in the estimated splice site usages between mutated and control samples in log scale. There is no RNA-seq read evidence supporting the novel splice sites for the control samples in genes IDUA, CORO1B, SHPRH, and ELOVL1, hence we report the usage estimate as 0 and the effect size for these genes as $\infty$.

### 2 Methods

In this section, we introduce the notations used to describe our model. Then, we introduce the inhomogeneous Markov chain model of McSplicer, the likelihood of the model's parameters, and the EM algorithm for estimation of parameters.

#### 2.1 Notations

We assume that potential exon start and end sites for a gene are given. This information can be obtained from known gene annotations or inferred from RNA-seq data using other methods. Suppose we have $M_s$ exon start sites, $s_1,\ldots,s_{M_s}$, and $M_e$ exon end sites $e_1,\ldots,e_{M_e}$. These $M_s$ and $M_e$ sites partition a gene into $M$ segments, $X_1,\ldots,X_M$, where $M = M_s + M_e + 1$. We introduce a sequence of hidden variables, $Z = (Z_1,\ldots,Z_M)$, where $Z_i$ is an indicator for whether a segment $X_i$ is a part of a transcript [1] ($Z_i = 1$) or not ($Z_i = 0$).

We define a subpath $s$ by a sequence of states for $(Z_a,\ldots,Z_b)$, $1 \le a \le b \le M$. Specifically, a subpath $s = z_{[a:b]}(o_a,\ldots,o_b)$, where $o_i \in \{0,1\}$ for $i = a,\ldots,b$, is defined by $Z_a = o_a, Z_{a+1} = o_{a+1},\ldots,Z_b = o_b$. In other words, a subpath $s = z_{[a:b]}(o_a,\ldots,o_b)$ describes whether each of the segments from $X_a$ to $X_b$ belongs to a transcript or not. Then, the probability of a subpath $s$ is:

$$\mathsf{P}(z_{[a:b]}(o_a,\ldots,o_b)) = \mathsf{P}(Z_a = o_a, Z_{a+1} = o_{a+1},\ldots,Z_b = o_b), \tag{1}$$

which is given by our inhomogeneous Markov chain model. A path $t$ is a subpath with $a = 1$ and $b = M$. A transcript can be represented by a path $t$, i.e., a sequence of states for $Z = (Z_1,\ldots,Z_M)$. Figure S13 shows an illustrative example of a gene with three transcripts which have four exon start sites and three exon end sites. These splice sites divide the gene into eight segments ($M_s = 4$, $M_e = 3$, and $M = 8$). For example, a path $t = z_{[1:8]}(1,1,0,1,0,0,1,1)$ (i.e., $Z = (1,1,0,1,0,0,1,1)$) indicates transcript $t_1$, and a subpath $s = z_{[3:5]}(0,1,0)$ indicates a subpath obtained from the same transcript; $t_1$.

We define the length of a subpath $s = z_{[a:b]}(o_a,\ldots,o_b)$, denoted by $l(s)$, by the number of bases included as a part of a transcript:

$$l(s) = l(z_{[a:b]}(o_a,\ldots,o_b)) = \sum_{a \le i \le b : o_i = 1} l(X_i), \tag{2}$$

where $l(X_j)$ is the number of bases in segment $X_j$. In the example of Figure S13, let's consider a subpath of the transcript $t_1$, $s = z_{[3:5]}(0,1,0)$. Then, $l(s) = l(z_{[3:5]}(1,0,1)) = l(X_3) + l(X_5)$. Similarly, we can define the length of a transcript (or a path), denoted by $l(t)$, by the number of bases included in the exonic regions of that transcript:

$$l(t) = l(z_{[1:M]}(o_1,\ldots,o_M)) = \sum_{1 \le i \le M : o_i = 1} l(X_i). \tag{3}$$

In the example of Figure S13, the transcript $t_1$ has the length $l(t_1) = l(z_{[1:8]}(1,1,0,1,0,0,1,1)) = l(X_1) + l(X_2) + l(X_4) + l(X_7) + l(X_8)$.

We use $F(s)$ to denote the index for the first segment in a subpath $s$ which is a part of a transcript, and use $L(s)$ to denote the index for the last segment in a subpath $s$ which is a part of a transcript. In the

[1] We use transcript and isoform interchangeably to refer to one splice variant of a gene. A gene usually has many splice variants.

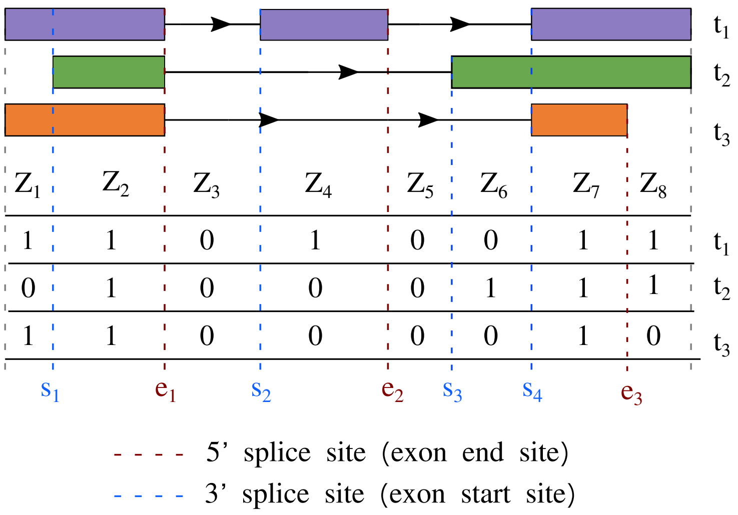


Figure S13: Hidden variables for segments defined by 3 different transcripts. The three sequences $(1,1,0,1,0,0,1,1), (0,1,0,0,0,1,1,1)$, and $(1,1,0,0,0,0,1,0)$ represent the three transcripts $t_1, t_2$, and $t_3$, respectively.

example of Figure S13, let's consider a subpath of $t_1$, $s = z_{[3:5]}(0,1,0)$. Then, $F(s) = 3$ and $L(s) = 5$. For the transcript $t_1$, the path $t_1 = z_{[1:8]}(1,1,0,1,0,0,1,1)$, $t_2(F) = 1$ and $t_2(L) = 8$. For the transcript $t_2$, the path $t_2 = z_{[1:8]}(0,1,0,1,0,1,1,1)$, $t_2(F) = 2$ and $t_2(L) = 8$. And similarly, For the transcript $t_3$, the path $t_3 = z_{[1:8]}(1,1,0,0,0,0,1,0)$, $t_3(F) = 1$ and $t_3(L) = 7$.

| transcript $t$ | $Z = (Z_1, \ldots, Z_8)$ | $w(t) = \mathsf{P}(Z_1, \ldots, Z_8)$ |
|---|---|---|
| $t_1$ | $z_{[1:8]}(1,1,0,1,0,0,1,1)$ | $\pi \times 1 \times q_1 \times p_2 \times q_2 \times (1-p_3) \times p_4 \times (1-q_3) \times q_4$ |
| $t_2$ | $z_{[1:8]}(0,1,0,0,0,1,1,1)$ | $(1-\pi) \times p_1 \times q_1 \times (1-p_2) \times 1 \times p_3 \times 1 \times (1-q_3) \times q_4$ |
| $t_3$ | $z_{[1:8]}(1,1,0,0,0,0,1,0)$ | $\pi \times 1 \times q_1 \times (1-p_2) \times 1 \times (1-p_3) \times p_4 \times q_3 \times 1$ |

Table S3: The relative abundances defined by the McSplicer model for the three transcripts presented in Fig. S13.

#### 2.2 An inhomogeneous Markov chain model

Now, we assume that $Z = (Z_1, \ldots, Z_M)$ follow an inhomogeneous Markov chain. Specifically, for the first segment $X_1$,

$$\mathsf{P}(Z_1 = 1) = \pi. \tag{4}$$

For two consecutive segments $X_i$ and $X_{i+1}$ for $i = 1, \ldots, M$, if they are separated by exon start site $s_m$ for $m = 1, \ldots, M_s$ (i.e., $i = I(s_m)$, where $I(s_m)$ is an index of the segment which appears on the left side of

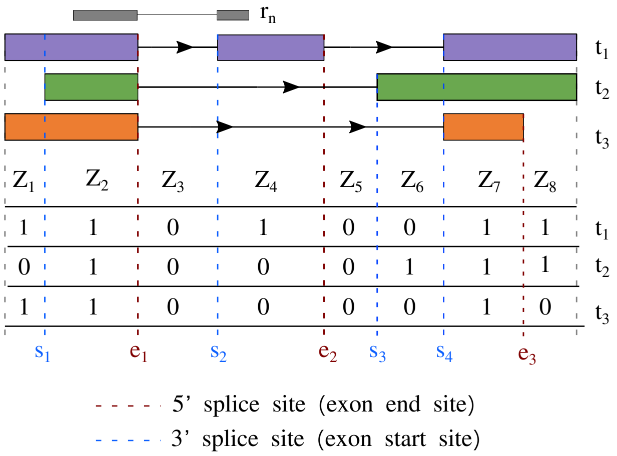


Figure S14: An example of a gene with three transcripts, the same as the one shown in Figure S13. Here, we show $r_n$, an observed sequence of read $n$, which was derived from the first transcript ($T_n = t_1$).

the exon start site $s_m$),

$$\mathsf{P}(Z_{i+1} = 1 | Z_i = 0) = p_m, \tag{5}$$

$$\mathsf{P}(Z_{i+1} = 1 | Z_i = 1) = 1. \tag{6}$$

If they are separated by exon end site $e_m$ for $m = 1, \ldots, M_e$ (i.e., $i = I(e_m)$, where $I(e_m)$ is an index of the segment which appears on the left side of the exon end site $e_m$),

$$\mathsf{P}(Z_{i+1} = 0 | Z_i = 0) = 1, \tag{7}$$

$$\mathsf{P}(Z_{i+1} = 0 | Z_i = 1) = q_m. \tag{8}$$

With this transition probability, we do not allow transcripts where $Z_i = 1$ and $Z_{i+1} = 0$ for $i = I(s_m)$, or $Z_i = 0$ and $Z_{i+1} = 1$ for $i = I(e_m)$. The parameters $p = (p_1, \ldots, p_{M_s})$ and $q = (q_1, \ldots, q_{M_e})$ indicate probabilities of using exon start sites and end sites, respectively. Precisely, these are conditional probabilities given that each site is considered for potential use. For example, with the current segment of being a part of a transcript (i.e., $Z_i = 1$), splicing process ignores exon start site without consideration (i.e., $\mathsf{P}(Z_{i+1} = 1 | Z_i = 1) = 1$ if $i = I(s_m)$) while it considers exon end site for potential use (i.e., $\mathsf{P}(Z_{i+1} = 0 | Z_i = 1) = q_m$ if $i = I(e_m)$). Table S3 lists probabilities for the three transcripts (or paths) in Figure S13 under our Markov model. Furthermore, to handle different transcript start and end sites within a gene, we introduce artificial starting and end points (i.e., reference points) in the implementation of this model.

#### 2.3 Likelihood of the parameters $\Theta = (\pi, p_1, \ldots, p_{M_s}, q_1, \ldots, q_{M_e})$

Following the notations introduced in [4]. Suppose we have RNA-seq reads mapped to a particular gene. The reads are derived from one end of each of the $N$ fragments and each read has length $L$. We assume that each fragment is independently generated from one of the possible transcripts allowed by our model. We denote the sequence of the $n$-th read as $R_n$. $T_n$ represents the transcript from which $R_n$ was generated. $B_n$ denotes the start position of $R_n$ in $T_n$.

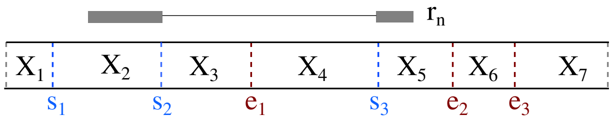


Figure S15: An example of a read which is not compatible with our model. The $S_n$ for $r_n$ in this example is $z_{[2:5]}(1,0,0,1)$. The two segments $X_2$ and $X_3$ are separated by the potential exon start site $s_2$. So our model doesn't allow $S_n = z_{[2:5]}(1,0,0,1)$ which implies $Z_2 = 1$ and $Z_3 = 0$. So $r_n$ is not compatible with our model.

For example, Figure S14 shows that $r_n$, an observed sequence of read $n$ was derived from the first transcript (i.e., $T_n = t_1$). So $T_n = z_{[1:8]}(1,1,0,1,0,0,1,1)$. The shortest subpath of $T_n$ from which read $n$ is derived, $S_n = z_{[2:4]}(1,0,1)$. Due to the definition of $S_n$, $F(S_n)$ is the index for the first segment in $S_n$, and $L(S_n)$ is the index for the last segment in $S_n$. In the example of Figure S14, for $S_n = z_{[2:4]}(1,0,1)$, $F(S_n) = 2$ and $L(S_n) = 4$.

To assume that all $r_n$ are derived from transcripts which are allowed in our model (i.e., $\mathsf{P}(r_n) > 0$ for all $r_n$), we filter out reads which are not compatible with our model. Figure S15 shows an example of a read which is not compatible with our model.

The likelihood of $\Theta$ can be written as

$$\mathsf{P}(r|\Theta) = \prod_{n=1}^{N} \mathsf{P}(r_n|\Theta) \tag{9}$$

$$= \prod_{n=1}^{N} [\sum_{t} \mathsf{P}(r_n, T_n = t|\Theta)] \tag{10}$$

$$= \prod_{n=1}^{N} [\sum_{t} [\sum_{(s,b):s\subset t} \mathsf{P}(r_n, S_n = s, B_n = b, T_n = t|\Theta)]] \tag{11}$$

$$\text{where } s \subset t \text{ means } s \text{ is a subpath of } t, \tag{12}$$

$$= \prod_{n=1}^{N} [\sum_{t} [\sum_{(s,b):s\subset t} \mathsf{P}(r_n|S_n = s, B_n = b)\mathsf{P}(S_n = s, B_n = b|T_n = t)\mathsf{P}(T_n = t|\Theta)]] \tag{13}$$

$$= \prod_{n=1}^{N} [\sum_{t} [\sum_{(s,b):s\subset t,(s,b)\to r_n} 1 \frac{1}{l(t)} \frac{l(t)w_\Theta(t)}{D(\Theta)}]] \tag{14}$$

$$\text{where } (s,b) \to r_n \text{ denotes that } r_n \text{ is the length } L \text{ sequence starting at position } b \tag{15}$$

$$\text{in the concatenation of segments in } s, \tag{16}$$

$$= \prod_{n=1}^{N} [\sum_{t} [\sum_{(s,b):s\subset t,(s,b)\to r_n} \frac{w_\Theta(t)}{D(\Theta)}]], \tag{17}$$

where $D(\Theta) = \sum_t l(t)w_\Theta(t)$. $l(t)$ represents the length of the transcript $t$, and $w_\Theta(t)$ represents the relative frequency (probability) of the transcript $t$.

##### 2.4 Parameter estimation

We use an EM algorithm to compute the maximum likelihood estimate for the model parameters $\Theta = \{\pi, p, q\}$, that is, $\hat{\Theta} := \text{argmax}_{\Theta}\mathsf{P}(r|\Theta)$. In this section, we first describe several useful quantities that are used in the EM algorithm, and then we explain our EM algorithm.

###### 2.4.1 Useful quantities

This section describes several useful quantities that are used in the EM algorithm. These quantities can be described as recursive form and computed efficiently using dynamic programming (DP).

###### 2.4.1.1 $\mathsf{P}(Z_i = 1)$ and $\mathsf{P}(Z_i = 0)$

We can compute $\mathsf{P}(Z_i = 1)$ the probability that segment $X_i$ is a part of a transcript and $\mathsf{P}(Z_i = 0)$ the probability that segment $X_i$ is not a part of a transcript, using DP as follows.

$$\begin{aligned}
\mathsf{P}(Z_i = 1) &= \mathsf{P}(Z_{i-1} = 1)\mathsf{P}(Z_i = 1|Z_{i-1} = 1) + \mathsf{P}(Z_{i-1} = 0)\mathsf{P}(Z_i = 1|Z_{i-1} = 0), &(18)\\
\mathsf{P}(Z_i = 0) &= 1 - \mathsf{P}(Z_i = 1), \text{ or equivalently} &(19)\\
&= \mathsf{P}(Z_{i-1} = 1)\mathsf{P}(Z_i = 0|Z_{i-1} = 1) + \mathsf{P}(Z_{i-1} = 0)\mathsf{P}(Z_i = 0|Z_{i-1} = 0), &(20)\\
\mathsf{P}(Z_1 = 0) &= 1 - \pi, &(21)\\
\mathsf{P}(Z_1 = 1) &= \pi. &(22)
\end{aligned}$$

If segments $X_{i-1}$ and $X_i$ are separated by exon start site $s_m$ (i.e., $i-1 = I(s_m)$),

$$\begin{aligned}
\mathsf{P}(Z_i = 1|Z_{i-1} = 0) &= p_m &(23)\\
\mathsf{P}(Z_i = 1|Z_{i-1} = 1) &= 1, &(24)\\
\mathsf{P}(Z_i = 0|Z_{i-1} = 0) &= 1 - p_m &(25)\\
\mathsf{P}(Z_i = 0|Z_{i-1} = 1) &= 0, &(26)
\end{aligned}$$

and if segments $X_{i-1}$ and $X_i$ are separated by exon end site $e_m$ (i.e., $i-1 = I(e_m)$),

$$\begin{aligned}
\mathsf{P}(Z_i = 1|Z_{i-1} = 0) &= 0 &(27)\\
\mathsf{P}(Z_i = 1|Z_{i-1} = 1) &= 1 - q_m, &(28)\\
\mathsf{P}(Z_i = 0|Z_{i-1} = 0) &= 1 &(29)\\
\mathsf{P}(Z_i = 0|Z_{i-1} = 1) &= q_m. &(30)
\end{aligned}$$

###### 2.4.1.2 $l_p(i, \mathbf{in})$, $l_p(i, \mathbf{out})$, $l_s(i, \mathbf{in})$, $l_s(i, \mathbf{out})$

- **Expected prefix length:** We define two types of the expected prefix length for the $i-th$ segment, $l_p(i, \text{in})$ and $l_p(i, \text{out})$, as follows. Let $Z_{[1:i]}$ denote a subpath which describes a sequence of states for $(Z_1, \ldots, Z_i)$. Then, the length of the subpath $Z_{[1:i]}$ is given by

$$l(Z_{[1:i]}) = \sum_{1 \le j \le i : Z_j = 1} l(X_j), \quad (31)$$

where $l(X_j)$ indicates the number of exonic bases in the segment $X_j$. $l_p(i,\text{in})$ is defined by the expected length of the subpath $Z_{[1:i]}$ given that $X_i$ is a part of a transcript (i.e., $Z_i = 1$) and $l_p(i,\text{out})$ is defined by the expected length of the subpath $Z_{[1:i]}$ given that $X_i$ is not a part of a transcript (i.e., $Z_i = 0$). Specifically,

$$l_p(i,\text{in}) = \mathsf{E}(l(Z_{[1:i]})|Z_i = 1) \tag{32}$$

$$l_p(i,\text{out}) = \mathsf{E}(l(Z_{[1:i]})|Z_i = 0). \tag{33}$$

We can compute these quantities using DP as follows.

$$l_p(i,\text{in}) = \mathsf{E}(l(Z_{[1:i]})|Z_i = 1) \tag{34}$$

$$= l(X_i) + \mathsf{E}(l(Z_{[1:(i-1)]}), Z_{i-1} = 1|Z_i = 1) + \mathsf{E}(l(Z_{[1:(i-1)]}), Z_{i-1} = 0|Z_i = 1) \tag{35}$$

$$= l(X_i) + \mathsf{E}(l(Z_{[1:(i-1)]})|Z_{i-1} = 1, Z_i = 1)\mathsf{P}(Z_{i-1} = 1|Z_i = 1) \tag{36}$$

$$+ \mathsf{E}(l(Z_{[1:(i-1)]})|Z_{i-1} = 0, Z_i = 1)\mathsf{P}(Z_{i-1} = 0|Z_i = 1) \tag{37}$$

$$\text{because conditional on } Z_{i-1}, Z_{[1:(i-1)]} \text{ and } Z_i \text{ are independent,} \tag{38}$$

$$= l(X_i) + \mathsf{E}(l(Z_{[1:(i-1)]})|Z_{i-1} = 1)\mathsf{P}(Z_{i-1} = 1|Z_i = 1) \tag{39}$$

$$+ \mathsf{E}(l(Z_{[1:(i-1)]})|Z_{i-1} = 0)\mathsf{P}(Z_{i-1} = 0|Z_i = 1) \tag{40}$$

$$= l(X_i) + l_p(i-1,\text{in})\frac{\mathsf{P}(Z_{i-1} = 1)\mathsf{P}(Z_i = 1|Z_{i-1} = 1)}{\mathsf{P}(Z_i = 1)} \tag{41}$$

$$+ l_p(i-1,\text{out})\frac{\mathsf{P}(Z_{i-1} = 0)\mathsf{P}(Z_i = 1|Z_{i-1} = 0)}{\mathsf{P}(Z_i = 1)}. \tag{42}$$

Similarly

$$l_p(i,\text{out}) = \mathsf{E}(l(Z_{[1:i]})|Z_i = 0) \tag{43}$$

$$= \mathsf{E}(l(Z_{[1:(i-1)]}), Z_{i-1} = 1|Z_i = 0) + \mathsf{E}(l(Z_{[1:(i-1)]}), Z_{i-1} = 0|Z_i = 0) \tag{44}$$

$$= l_p(i-1,\text{in})\frac{\mathsf{P}(Z_{i-1} = 1)\mathsf{P}(Z_i = 0|Z_{i-1} = 1)}{\mathsf{P}(Z_i = 0)} \tag{45}$$

$$+ l_p(i-1,\text{out})\frac{\mathsf{P}(Z_{i-1} = 0)\mathsf{P}(Z_i = 0|Z_{i-1} = 0)}{\mathsf{P}(Z_i = 0)}. \tag{46}$$

$$\tag{47}$$

And,

$$l_p(1,\text{in}) = l(X_1) \tag{48}$$

$$l_p(1,\text{out}) = 0. \tag{49}$$

Here, $\mathsf{P}(Z_{i-1} = 0), \mathsf{P}(Z_{i-1} = 1), \mathsf{P}(Z_i = 0), \mathsf{P}(Z_i = 1), \mathsf{P}(Z_i = 0|Z_{i-1} = 1), \mathsf{P}(Z_i = 1|Z_{i-1} = 1), \mathsf{P}(Z_i = 0|Z_{i-1} = 0), \mathsf{P}(Z_i = 1|Z_{i-1} = 0)$ can be computed as described in Section 2.4.1.1.

- **Expected suffix length:** We define two types of the expected suffix length for the $i-th$ segment , $l_s(i,\text{in})$ and $l_s(i,\text{out})$, as follows. Let $Z_{[i:M]}$ denote a subpath which describes a sequence of states for $(Z_i,\dots,Z_M)$. $l_s(i,\text{in})$ is defined by the expected length of the subpath $Z_{[i:M]}$ given that $X_i$ is a part of an isoform (i.e., $Z_i = 1$) and $l_s(i,\text{out})$ is defined by the expected length of the subpath $Z_{[i:M]}$ given that $X_i$ is not a part of an isoform (i.e., $Z_i = 0$). Specifically,

$$l_s(i,\text{in}) = \mathsf{E}(l(Z_{[i:M]})|Z_i = 1) \tag{50}$$
$$l_s(i,\text{out}) = \mathsf{E}(l(Z_{[i:M]})|Z_i = 0). \tag{51}$$

We can compute these quantities using DP as follows.

$$l_s(i,\text{in}) = \mathsf{E}(l(Z_{[i:M]})|Z_i = 1) \tag{52}$$
$$= l(X_i) + \mathsf{E}(l(Z_{[(i+1):M]}), Z_{i+1} = 1|Z_i = 1) + \mathsf{E}(l(Z_{[(i+1):M]}), Z_{i+1} = 0|Z_i = 1) \tag{53}$$
$$= l(X_i) + \mathsf{E}(l(Z_{[(i+1):M]})|Z_{i+1} = 1, Z_i = 1)\mathsf{P}(Z_{i+1} = 1|Z_i = 1) \tag{54}$$
$$+ \mathsf{E}(l(Z_{[(i+1):M]})|Z_{i+1} = 0, Z_i = 1)\mathsf{P}(Z_{i+1} = 0|Z_i = 1) \tag{55}$$
$$\text{because conditional on } Z_{i+1}, Z_{[(i+1):M]} \text{ and } Z_i \text{ are independent,} \tag{56}$$
$$= l(X_i) + \mathsf{E}(l(Z_{[(i+1):M]})|Z_{i+1} = 1)\mathsf{P}(Z_{i+1} = 1|Z_i = 1) \tag{57}$$
$$+ \mathsf{E}(l(Z_{[(i+1):M]})|Z_{i+1} = 0)\mathsf{P}(Z_{i+1} = 0|Z_i = 1) \tag{58}$$
$$= l(X_i) + l_s(i+1,\text{in})\mathsf{P}(Z_{i+1} = 1|Z_i = 1) \tag{59}$$
$$+ l_s(i+1,\text{out})\mathsf{P}(Z_{i+1} = 0|Z_i = 1). \tag{60}$$

Similarly

$$l_s(i,\text{out}) = \mathsf{E}(l(Z_{[i:M]})|Z_i = 0) \tag{61}$$
$$= \mathsf{E}(l(Z_{[(i+1):M]}), Z_{i+1} = 1|Z_i = 0) + \mathsf{E}(l(Z_{[(i+1):M]}), Z_{i+1} = 0|Z_i = 0) \tag{62}$$
$$= \mathsf{E}(l(Z_{[(i+1):M]})|Z_{i+1} = 1, Z_i = 0)\mathsf{P}(Z_{i+1} = 1|Z_i = 0) \tag{63}$$
$$+ \mathsf{E}(l(Z_{[(i+1):M]})|Z_{i+1} = 0, Z_i = 0)\mathsf{P}(Z_{i+1} = 0|Z_i = 0) \tag{64}$$
$$= l_s(i+1,\text{in})\mathsf{P}(Z_{i+1} = 1|Z_i = 0) \tag{65}$$
$$+ l_s(i+1,\text{out})\mathsf{P}(Z_{i+1} = 0|Z_i = 0). \tag{66}$$

And,

$$l_s(M,\text{in}) = l(X_M) \tag{67}$$
$$l_s(M,\text{out}) = 0. \tag{68}$$

Here, $\mathsf{P}(Z_{i+1} = 0|Z_i = 1), \mathsf{P}(Z_{i+1} = 1|Z_i = 1), \mathsf{P}(Z_{i+1} = 0|Z_i = 0), \mathsf{P}(Z_{i+1} = 1|Z_i = 0)$ can be computed as described in Section 2.4.1.1

###### 2.4.1.3 $f_{11}(i,j)$, $f_{10}(i,j)$, $f_{01}(i,j)$, and $f_{00}(i,j)$

For $1 \le i \le j \le M$, $f_{**}(i,j)$ is defined by the probability of $Z_j$ conditional on $Z_i$. Specifically,

$$f_{11}(i,j) := \mathsf{P}(Z_j = 1|Z_i = 1) \tag{69}$$
$$f_{10}(i,j) := \mathsf{P}(Z_j = 0|Z_i = 1) \tag{70}$$
$$f_{01}(i,j) := \mathsf{P}(Z_j = 1|Z_i = 0) \tag{71}$$
$$f_{00}(i,j) := \mathsf{P}(Z_j = 0|Z_i = 0). \tag{72}$$

We can compute these quantities using DP as follows.

- **When** $i = j$

$$f_{11}(i,j) := \mathsf{P}(Z_j = 1|Z_i = 1) = 1 \tag{73}$$
$$f_{10}(i,j) := \mathsf{P}(Z_j = 0|Z_i = 1) = 0 \tag{74}$$
$$f_{01}(i,j) := \mathsf{P}(Z_j = 1|Z_i = 0) = 0 \tag{75}$$
$$f_{00}(i,j) := \mathsf{P}(Z_j = 0|Z_i = 0) = 1. \tag{76}$$

- **When** $i < j$
**If two segments** $X_{j-1}$ **and** $X_j$ **are separated by potential exon start site** $s_m$ **(i.e.,** $j-1 = I(s_m)$**):**

$$f_{11}(i,j) := \mathsf{P}(Z_j = 1|Z_i = 1) = \begin{cases} 1 \text{ if } i = j-1 \\ f_{11}(i,j-1) + f_{10}(i,j-1)p_m \text{ if } i < j-1, \end{cases} \tag{77}$$

because $\mathsf{P}(Z_j = 1|Z_i = 1)$

$$= \mathsf{P}(Z_j = 1, Z_{j-1} = 1|Z_i = 1) + \mathsf{P}(Z_j = 1, Z_{j-1} = 0|Z_i = 1) \tag{78}$$
$$= \mathsf{P}(Z_j = 1|Z_{j-1} = 1, Z_i = 1)\mathsf{P}(Z_{j-1} = 1|Z_i = 1) + \mathsf{P}(Z_j = 1|Z_{j-1} = 0, Z_i = 1)\mathsf{P}(Z_{j-1} = 0|Z_i = 1) \tag{79}$$
$$= \mathsf{P}(Z_j = 1|Z_{j-1} = 1)\mathsf{P}(Z_{j-1} = 1|Z_i = 1) + \mathsf{P}(Z_j = 1|Z_{j-1} = 0)\mathsf{P}(Z_{j-1} = 0|Z_i = 1) \tag{80}$$
$$= f_{11}(i,j-1) + p_m f_{10}(i,j-1). \tag{81}$$

Similarly,

$$f_{10}(i,j) := \mathsf{P}(Z_j = 0|Z_i = 1) = \begin{cases} 0 & \text{if } i = j-1 \\ f_{10}(i,j-1)(1-p_m) & \text{if } i < j-1. \end{cases} \tag{82}$$

$$f_{01}(i,j) := \mathsf{P}(Z_j = 1|Z_i = 0) = \begin{cases} p_m & \text{if } i = j-1 \\ f_{01}(i,j-1) + f_{00}(i,j-1)p_m & \text{if } i < j-1. \end{cases} \tag{83}$$

$$f_{00}(i,j) := \mathsf{P}(Z_j = 0|Z_i = 0) = \begin{cases} 1-p_m & \text{if } i = j-1 \\ f_{00}(i,j-1)(1-p_m) & \text{if } i < j-1. \end{cases} \tag{84}$$

**If two segments** $X_{j-1}$ **and** $X_j$ **are separated by potential exon end site** $e_m$ **(i.e.,** $j-1 = I(e_m)$**):**

$$f_{11}(i,j) := \mathsf{P}(Z_j = 1|Z_i = 1) = \begin{cases} 1-q_m & \text{if } i = j-1 \\ f_{11}(i,j-1)(1-q_m) & \text{if } i < j-1. \end{cases} \tag{85}$$

$$f_{10}(i,j) := \mathsf{P}(Z_j = 0|Z_i = 1) = \begin{cases} q_m & \text{if } i = j-1 \\ f_{10}(i,j-1) + f_{11}(i,j-1)q_m & \text{if } i < j-1. \end{cases} \tag{86}$$

$$f_{01}(i,j) := \mathsf{P}(Z_j = 1|Z_i = 0) = \begin{cases} 0 & \text{if } i = j-1 \\ f_{01}(i,j-1)(1-q_m) & \text{if } i < j-1. \end{cases} \tag{87}$$

$$f_{00}(i,j) := \mathsf{P}(Z_j = 0|Z_i = 0) = \begin{cases} 1 & \text{if } i = j-1 \\ f_{00}(i,j-1) + f_{01}(i,j-1)q_m & \text{if } i < j-1. \end{cases} \tag{88}$$

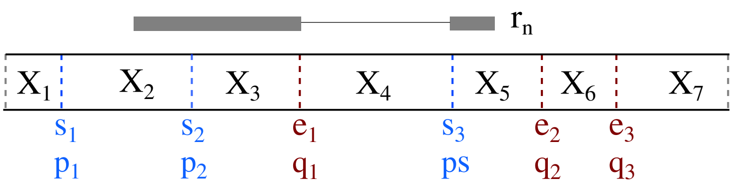


Figure S16: In this example, $S_n = z_{[2:5]}(1,1,0,1)$ and $w(s) = \mathsf{P}(Z_3 = 1, Z_4 = 0, Z_5 = 1 | Z_2 = 1) = 1 \cdot q_1 \cdot p_3$

###### 2.4.1.4 $w(s) = \frac{\mathsf{P}(S_n = s)}{\mathsf{P}(Z_{F(s)} = 1)} = \mathsf{P}(S_n = s | Z_{F(s)} = 1)$

We use $w(s)$ to denote the conditional probability of $S_n = s$ conditional on $X_{F(s)}$ is a part of an isoform. Let a subpath $s = z_{[a:b]}(o_a, \ldots, o_b)$. Due to the definition of $S_n$, that is the shortest subpath of $T_n$ from which read $n$ is derived, $o_a = 1, o_b = 1, F(s) = a$ and $L(s) = b$.

$$w(s) = \frac{\mathsf{P}(S_n = s)}{\mathsf{P}(Z_{F(s)} = 1)} \tag{89}$$

$$= \frac{\mathsf{P}(Z_a = o_a, Z_{a+1} = o_{a+1}, \ldots, Z_b = o_b)}{\mathsf{P}(Z_a = 1)} \tag{90}$$

$$= \mathsf{P}(Z_{a+1} = o_{a+1}, \ldots, Z_b = o_b | Z_a = 1) \tag{91}$$

$$= \mathsf{P}(Z_a = 1, Z_{a+1} = o_{a+1}, \ldots, Z_b = o_b | Z_a = 1) \tag{92}$$

$$= \mathsf{P}(S_n = s | Z_{F(s)} = 1). \tag{93}$$

Moreover,

$$w(s) = \mathsf{P}(Z_{a+1} = o_{a+1}, \ldots, Z_b = o_b | Z_a = 1) \tag{94}$$

$$= \prod_{i=a}^{b-1} \mathsf{P}(Z_{i+1} = o_{i+1} | Z_i = o_i) \tag{95}$$

$$= \prod_{s_m : a \le I(s_m) < b} p_m^{(1-o_{I(s_m)})(o_{[I(s_m)+1]})} (1 - p_m)^{(1-o_{I(s_m)})(1-o_{[I(s_m)+1]})} \tag{96}$$

$$\times \prod_{e_m : a \le I(e_m) < b} q_m^{(o_{I(e_m)})(1-o_{[I(e_m)+1]})} (1 - q_m)^{(o_{I(e_m)})(o_{[I(e_m)+1]})}. \tag{97}$$

In the example of Figure S16, $s = z_{[2:5]}(1,1,0,1)$. So $a = 2, b = 5, o_a = 1, o_b = 1, F(s) = 2$ and

$L(s) = 5$. In this example,

$$
\begin{aligned}
w(s) &= \mathsf{P}(Z_3 = 1, Z_4 = 0, Z_5 = 1 | Z_2 = 1) && (98)\\
&= \mathsf{P}(Z_3 = 1 | Z_2 = 1)\mathsf{P}(Z_4 = 0 | Z_3 = 1)\mathsf{P}(Z_5 = 1 | Z_4 = 1) && (99)\\
&= 1 \cdot q_1 \cdot p_3 && (100)\\
&\text{or,} && (101)\\
&= \prod_{s_m : 2 \le I(s_m) < 5} p_m^{(1-o_{I(s_m)})(o_{[I(s_m)+1]})} (1-p_m)^{(1-o_{I(s_m)})(1-o_{[I(s_m)+1]})} && (102)\\
&\times \prod_{e_m : 2 \le I(e_m) < 5} q_m^{(o_{I(e_m)})(1-o_{[I(e_m)+1]})} (1-q_m)^{(o_{I(e_m)})(o_{[I(e_m)+1]})} && (103)\\
&= \prod_{s_m : s_2, s_3} p_m^{(1-o_{I(s_m)})(o_{[I(s_m)+1]})} (1-p_m)^{(1-o_{I(s_m)})(1-o_{[I(s_m)+1]})} && (104)\\
&\times \prod_{e_m : e_1} q_m^{(o_{I(e_m)})(1-o_{[I(e_m)+1]})} (1-q_m)^{(o_{I(e_m)})(o_{[I(e_m)+1]})} && (105)\\
&= p_2^{(1-1)(1)} (1-p_2)^{(1-1)(1-1)} p_3^{(1-0)(1)} (1-p_3)^{(1-0)(1-1)} && (106)\\
&\times q_1^{(1)(1-0)} (1-q_1)^{(1)(0)} && (107)\\
&= 1 \cdot p_3 \times q_1. && (108)
\end{aligned}
$$

#### 2.4.1.5 $w(s_l, s_r)$

Given paths $s_l$ and $s_r$ with $F(s_r) \ge F(s_l)$. Let $a := F(s_l)$, $b := L(s_l)$, $a' := F(s_r)$, $b' := L(s_r)$, and $m := a' - b - 1$. The probability $w_p(s_l, s_r) := \mathsf{P}(s_l \in S_n, s_r \in S_n | Z_a = 1)$ of read $n$ coming from a fragment starting with $s_l$ and ending with $s_r$, conditional on $X_a$ being part of the isoform, is $w(s_l \cdot s_r) f(l(s_l \cdot s_r))$ if $b \le a + 1$, and otherwise

$$
\begin{aligned}
w_p(s_l, s_r) &= \sum_{o_{b+1},\ldots,o_{a'-1} \in \{0,1\}^m} \mathsf{P}(S_n = s_l \cdot z_{[b+1,a'-1]}(o_{b+1},\ldots,o_{a'-1}) \cdot s_r | Z_a = 1) && (109)\\
&= \sum_{o_{b+1},\ldots,o_{a'-1} \in \{0,1\}^m} w(s_l \cdot z_{[b+1,a'-1]}(o_{b+1},\ldots,o_{a'-1}) \cdot s_r) f(l(s_l \cdot z_{[b+1:a'-1]}(o_{b+1},\ldots,o_{a'-1}) \cdot s_r)) && (110)
\end{aligned}
$$

We also define probability $w'_p(s_l, s_r, z_{[i:i+1]}(o_i, o_{i+1}))$ which restricts paths connecting $s_l$ and $s_r$ to use or not use a given start or end segment $i$:

$$
\begin{aligned}
& w_p(s_l, s_r, z_{[i:i+1]}(o'_i, o'_{i+1}))) = && (111)\\
& \sum_{\substack{o_{b+1},\ldots,o_{a'-1} \in \{0,1\}^m : \\ o_i = o'_i, o_{i+1} = o'_{i+1}}} w(s_l \cdot z_{[b+1,a'-1]}(o_{b+1},\ldots,o_{a'-1}) \cdot s_r) f(l(s_l \cdot z_{[b+1:a'-1]}(o_{b+1},\ldots,o_{a'-1}) \cdot s_r)) && (112)
\end{aligned}
$$

###### 2.4.1.6 $\mathsf{P}(Z^n_{I(s_m)} = 0, Z^n_{I(s_m)+1} = 1, S_n = S)$, $\mathsf{P}(Z^n_{I(s_m)} = 0, Z^n_{I(s_m)+1} = 0, S_n = S)$, $\mathsf{P}(Z^n_{I(e_m)} = 1, Z^n_{I(e_m)+1} = 0, S_n = S)$, and $\mathsf{P}(Z^n_{I(e_m)} = 1, Z^n_{I(e_m)+1} = 1, S_n = S)$

- $\mathsf{P}(Z^n_{I(s_m)} = 0, Z^n_{I(s_m)+1} = 1, S_n = s)$

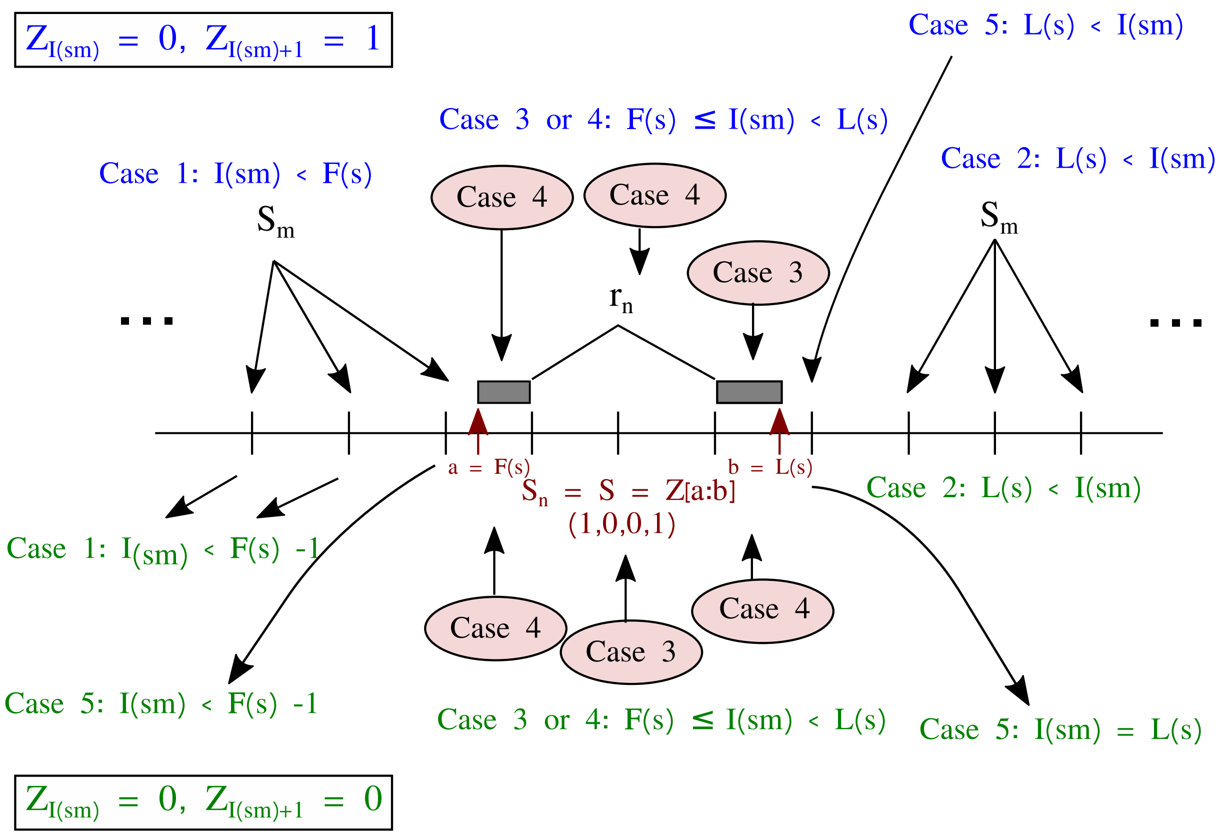


Figure S17: Visualize five cases considered for computations of $\mathsf{P}(Z^n_{I(s_m)}=0, Z^n_{I(s_m)+1}=1, S_n=s)$ and $\mathsf{P}(Z^n_{I(s_m)}=0, Z^n_{I(s_m)+1}=0, S_n=s)$

**case 1** $I(s_m) < F(s)$**:**
As shown in Figure S17, a potential exon start site $s_m$ appears left side of a subpath $s = z_{[a:b]}(o_a, \ldots, o_b)$.

$$\mathsf{P}(Z_{I(s_m)}=0, Z_{I(s_m)+1}=1, s) \tag{113}$$
$$= \mathsf{P}(Z_1=0, Z_{I(s_m)}=0, Z_{I(s_m)+1}=1, s) + \mathsf{P}(Z_1=1, Z_{I(s_m)}=0, Z_{I(s_m)+1}=1, s) \tag{114}$$
$$= \mathsf{P}(Z_1=0)\mathsf{P}(Z_{I(s_m)}=0|Z_1=0)\mathsf{P}(Z_{I(s_m)+1}=1|Z_{I(s_m)}=0)\mathsf{P}(Z_{F(s)}=1|Z_{I(s_m)+1}=1)\mathsf{P}(s|Z_{F(s)}=1) \tag{115}$$
$$+ \mathsf{P}(Z_1=1)\mathsf{P}(Z_{I(s_m)}=0|Z_1=1)\mathsf{P}(Z_{I(s_m)+1}=1|Z_{I(s_m)}=0)\mathsf{P}(Z_{F(s)}=1|Z_{I(s_m)+1}=1)\mathsf{P}(s|Z_{F(s)}=1) \tag{116}$$
$$= (1-\pi)f_{00}(1, I(s_m))p_m f_{11}(I(s_m)+1, F(s))w(s) + \pi f_{10}(1, I(s_m))p_m f_{11}(I(s_m)+1, F(s))w(s) \tag{117}$$
$$= \left[(1-\pi)f_{00}(1, I(s_m)) + \pi f_{10}(1, I(s_m))\right] \times p_m f_{11}(I(s_m)+1, F(s))w(s). \tag{118}$$

Here, $f_{**}(i,j)$ and $w(s)$ can be computed as described in Sections 2.4.1.3 2.4.1.4.

**case 2** $L(s) < I(s_m)$**:**

As shown in Figure S17, a potential exon start site $s_m$ appears right side of a subpath $s = z_{[a:b]}(o_a, \ldots, o_b)$.

$$\mathsf{P}(s, Z_{I(s_m)} = 0, Z_{I(s_m)+1} = 1) \tag{119}$$
$$= \mathsf{P}(Z_1 = 0, s, Z_{I(s_m)} = 0, Z_{I(s_m)+1} = 1) + \mathsf{P}(Z_1 = 1, s, Z_{I(s_m)} = 0, Z_{I(s_m)+1} = 1) \tag{120}$$
$$= \mathsf{P}(Z_1 = 0)\mathsf{P}(Z_{F(s)} = 1|Z_1 = 0)\mathsf{P}(s|Z_{F(s)} = 1)\mathsf{P}(Z_{I(s_m)} = 0|Z_{L(s)} = 1)\mathsf{P}(Z_{I(s_m)+1} = 1|Z_{I(s_m)} = 0) \tag{121}$$
$$+ \mathsf{P}(Z_1 = 1)\mathsf{P}(Z_{F(s)} = 1|Z_1 = 1)\mathsf{P}(s|Z_{F(s)} = 1)\mathsf{P}(Z_{I(s_m)} = 0|Z_{L(s)} = 1)\mathsf{P}(Z_{I(s_m)+1} = 1|Z_{I(s_m)} = 0) \tag{122}$$
$$= (1-\pi)f_{01}(1, F(s))w(s)f_{10}(L(s), I(s_m))p_m + \pi f_{11}(1, F(s))w(s)f_{10}(L(s), I(s_m))p_m \tag{123}$$
$$= \big[(1-\pi)f_{01}(1, F(s)) + \pi f_{11}(1, F(s))\big] \times w(s)f_{10}(L(s), I(s_m))p_m. \tag{124}$$

**case 3** $F(s) \le I(s_m) < L(s)$ **and** $(Z_{I(s_m)} = 0, Z_{I(s_m)+1} = 1)$ **is a subset of** $s$**:**
A subpath $s = z_{[a:b]}(o_a, \ldots, o_b)$ can be represented by $(Z_a = o_a, Z_{a+1} = o_{a+1}, \ldots, Z_b = o_b)$. If the subpath $s$ contains$(Z_{I(s_m)} = 0, Z_{I(s_m)+1} = 1)$, then $(Z_{I(s_m)} = 0, Z_{I(s_m)+1} = 1)$ is a subset of $s$ (i.e., $(Z_{I(s_m)} = 0, Z_{I(s_m)+1} = 1) \subset s$). As shown in Figure S17, a potential exon start site $s_m$ appears inside of a subpath $s = z_{[a:b]}(o_a, \ldots, o_b)$.

$$\mathsf{P}(s, Z_{I(s_m)} = 0, Z_{I(s_m)+1} = 1) \tag{125}$$
$$= \mathsf{P}(s) \tag{126}$$
$$= \mathsf{P}(Z_1 = 0, s) + \mathsf{P}(Z_1 = 1, s) \tag{127}$$
$$= \mathsf{P}(Z_1 = 0)\mathsf{P}(Z_{F(s)} = 1|Z_1 = 0)\mathsf{P}(s|Z_{F(s)} = 1) + \mathsf{P}(Z_1 = 1)\mathsf{P}(Z_{F(s)} = 1|Z_1 = 1)\mathsf{P}(s|Z_{F(s)} = 1) \tag{128}$$
$$= (1-\pi)f_{01}(1, F(s))w(s) + \pi f_{11}(1, F(s))w(s) \tag{129}$$
$$= \big[(1-\pi)f_{01}(1, F(s)) + \pi f_{11}(1, F(s))\big] \times w(s). \tag{130}$$

**case 4** $F(s) \le I(s_m) < L(s)$ **and** $(Z_{I(s_m)} = 0, Z_{I(s_m)+1} = 1)$ **is not a subset of** $s$**:**
This event is now allowed.

$$\mathsf{P}(s, Z_{I(s_m)} = 0, Z_{I(s_m)+1} = 1) = 0 \tag{131}$$

**case 5** $I(s_m) = L(s)$**:**
This event is now allowed.

$$\mathsf{P}(s, Z_{I(s_m)} = 0, Z_{I(s_m)+1} = 1) = 0 \tag{132}$$

- $\mathsf{P}(Z^n_{I(s_m)} = 0, Z^n_{I(s_m)+1} = 0, S_n = s)$
  **case 1** $I(s_m) < F(s) - 1$**:**

As shown in Figure S17, a potential exon start site $s_m$ appears left side of a subpath $s = z_{[a:b]}(o_a, \ldots, o_b)$.

$$\begin{aligned}
&\mathsf{P}(Z_{I(s_m)} = 0, Z_{I(s_m)+1} = 0, s) && (133)\\
&= \mathsf{P}(Z_1 = 0, Z_{I(s_m)} = 0, Z_{I(s_m)+1} = 0, s) + \mathsf{P}(Z_1 = 1, Z_{I(s_m)} = 0, Z_{I(s_m)+1} = 0, s) && (134)\\
&= \mathsf{P}(Z_1 = 0)\mathsf{P}(Z_{I(s_m)} = 0|Z_1 = 0)\mathsf{P}(Z_{I(s_m)+1} = 0|Z_{I(s_m)} = 0)\mathsf{P}(Z_{F(s)} = 1|Z_{I(s_m)+1} = 0)\mathsf{P}(s|Z_{F(s)} = 1) && (135)\\
&+ \mathsf{P}(Z_1 = 1)\mathsf{P}(Z_{I(s_m)} = 0|Z_1 = 1)\mathsf{P}(Z_{I(s_m)+1} = 0|Z_{I(s_m)} = 0)\mathsf{P}(Z_{F(s)} = 1|Z_{I(s_m)+1} = 0)\mathsf{P}(s|Z_{F(s)} = 1) && (136)\\
&= (1-\pi)f_{00}(1, I(s_m))(1-p_m)f_{01}(I(s_m)+1, F(s))w(s) + \pi f_{10}(1, I(s_m))(1-p_m)f_{01}(I(s_m)+1, F(s))w(s) && (137)\\
&= \left[(1-\pi)f_{00}(1, I(s_m)) + \pi f_{10}(1, I(s_m))\right] \times (1-p_m)f_{01}(I(s_m)+1, F(s))w(s). && (138)
\end{aligned}$$

**case 2 $L(s) < I(s_m)$:**
As shown in Figure S17, a potential exon start site $s_m$ appears right side of a subpath $s = z_{[a:b]}(o_a, \ldots, o_b)$.

$$\begin{aligned}
&\mathsf{P}(s, Z_{I(s_m)} = 0, Z_{I(s_m)+1} = 0) && (139)\\
&= \mathsf{P}(Z_1 = 0, s, Z_{I(s_m)} = 0, Z_{I(s_m)+1} = 0) + \mathsf{P}(Z_1 = 1, s, Z_{I(s_m)} = 0, Z_{I(s_m)+1} = 0) && (140)\\
&= \mathsf{P}(Z_1 = 0)\mathsf{P}(Z_{F(s)} = 1|Z_1 = 0)\mathsf{P}(s|Z_{F(s)} = 1)\mathsf{P}(Z_{I(s_m)} = 0|Z_{L(s)} = 1)\mathsf{P}(Z_{I(s_m)+1} = 0|Z_{I(s_m)} = 0) && (141)\\
&+ \mathsf{P}(Z_1 = 1)\mathsf{P}(Z_{F(s)} = 1|Z_1 = 1)\mathsf{P}(s|Z_{F(s)} = 1)\mathsf{P}(Z_{I(s_m)} = 0|Z_{L(s)} = 1)\mathsf{P}(Z_{I(s_m)+1} = 0|Z_{I(s_m)} = 0) && (142)\\
&= (1-\pi)f_{01}(1, F(s))w(s)f_{10}(L(s), I(s_m))(1-p_m) + \pi f_{11}(1, F(s))w(s)f_{10}(L(s), I(s_m))(1-p_m) && (143)\\
&= \left[(1-\pi)f_{01}(1, F(s)) + \pi f_{11}(1, F(s))\right] \times w(s)f_{10}(L(s), I(s_m))(1-p_m). && (144)
\end{aligned}$$

**case 3 $F(s) \leq I(s_m) < L(s)$ and $(Z_{I(s_m)} = 0, Z_{I(s_m)+1} = 0)$ is a subset of $s$:**
As shown in Figure S17, a potential exon start site $s_m$ appears inside of a subpath $s = z_{[a:b]}(o_a, \ldots, o_b)$.

$$\begin{aligned}
&\mathsf{P}(s, Z_{I(s_m)} = 0, Z_{I(s_m)+0} = 1) && (145)\\
&= \mathsf{P}(s) && (146)\\
&= \mathsf{P}(Z_1 = 0, s) + \mathsf{P}(Z_1 = 1, s) && (147)\\
&= \mathsf{P}(Z_1 = 0)\mathsf{P}(Z_{F(s)} = 1|Z_1 = 0)\mathsf{P}(s|Z_{F(s)} = 1) + \mathsf{P}(Z_1 = 1)\mathsf{P}(Z_{F(s)} = 1|Z_1 = 1)\mathsf{P}(s|Z_{F(s)} = 1) && (148)\\
&= (1-\pi)f_{01}(1, F(s))w(s) + \pi f_{11}(1, F(s))w(s) && (149)\\
&= \left[(1-\pi)f_{01}(1, F(s)) + \pi f_{11}(1, F(s))\right] \times w(s). && (150)
\end{aligned}$$

**case 4 $F(s) \leq I(s_m) < L(s)$ and $(Z_{I(s_m)} = 0, Z_{I(s_m)+1} = 0)$ is not a subset of $s$:**
This event is now allowed.

$$\mathsf{P}(s, Z_{I(s_m)} = 0, Z_{I(s_m)+1} = 0) = 0 \qquad (151)$$

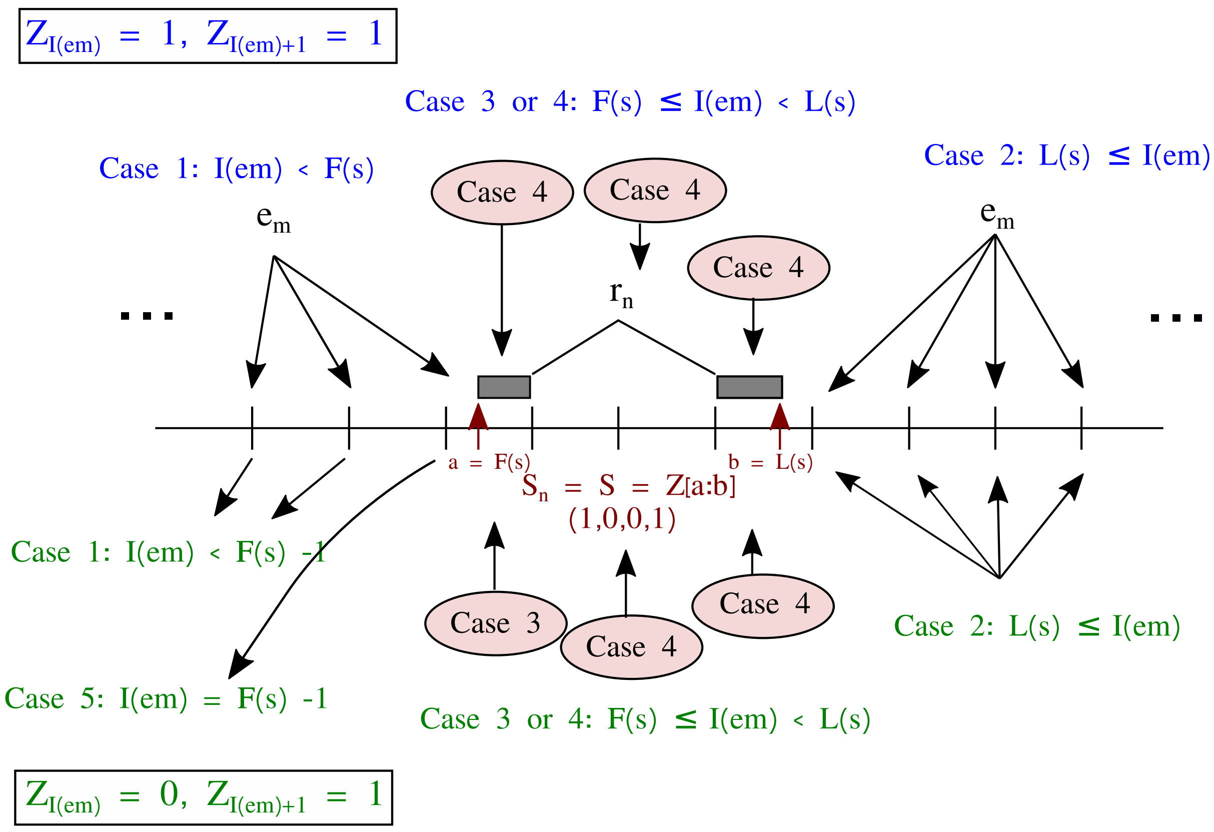


Figure S18: Visualize five cases considered for computations of $\mathsf{P}(Z^n_{I(e_m)} = 1, Z^n_{I(e_m)+1} = 0, S_n = s)$ and $\mathsf{P}(Z^n_{I(e_m)} = 1, Z^n_{I(e_m)+1} = 1, S_n = s)$

**case 5** $I(s_m) = F(s) - 1$ **or** $I(s_m) = L(s)$**:**
This event is now allowed.

$$\mathsf{P}(s, Z_{I(s_m)} = 0, Z_{I(s_m)+1} = 0) = 0 \tag{152}$$

- $\mathsf{P}(Z^n_{I(e_m)} = 1, Z^n_{I(e_m)+1} = 1, S_n = s)$
  **case 1** $I(e_m) < F(s)$**:**
  As shown in Figure S18, a potential exon end site $e_m$ appears left side of a subpath $s = z_{[a:b]}(o_a, \ldots, o_b)$.

$$\mathsf{P}(Z_{I(e_m)} = 1, Z_{I(e_m)+1} = 1, s) \tag{153}$$

$$= \mathsf{P}(Z_1 = 0, Z_{I(e_m)} = 1, Z_{I(e_m)+1} = 1, s) + \mathsf{P}(Z_1 = 1, Z_{I(e_m)} = 1, Z_{I(e_m)+1} = 1, s) \tag{154}$$

$$= (1-\pi) f_{01}(1, I(e_m))(1-q_m) f_{11}(I(e_m)+1, F(s)) w(s) + \pi f_{11}(1, I(e_m))(1-q_m) f_{11}(I(e_m)+1, F(s)) w(s) \tag{155}$$

$$= \left[(1-\pi) f_{01}(1, I(e_m)) + \pi f_{11}(1, I(e_m))\right] \times (1-q_m) f_{11}(I(e_m)+1, F(s)) w(s). \tag{156}$$

**case 2** $L(s) \le I(s_m)$**:**
As shown in Figure S18, a potential exon end site $e_m$ appears right side of a subpath $s = z_{[a:b]}(o_a, \ldots, o_b)$.

$$\mathsf{P}(s, Z_{I(e_m)} = 1, Z_{I(e_m)+1} = 1) \tag{157}$$
$$= \mathsf{P}(Z_1 = 0, s, Z_{I(e_m)} = 1, Z_{I(e_m)+1} = 1) + \mathsf{P}(Z_1 = 1, s, Z_{I(e_m)} = 1, Z_{I(e_m)+1} = 1) \tag{158}$$
$$= (1-\pi) f_{01}(1, F(s)) w(s) f_{11}(L(s), I(e_m))(1-q_m) + \pi f_{11}(1, F(s)) w(s) f_{11}(L(s), I(e_m))(1-q_m) \tag{159}$$
$$= \big[(1-\pi) f_{01}(1, F(s)) + \pi f_{11}(1, F(s))\big] \times w(s) f_{11}(L(s), I(e_m))(1-q_m). \tag{160}$$

**case 3** $F(s) \le I(e_m) < L(s)$ **and** $(Z_{I(e_m)} = 1, Z_{I(e_m)+1} = 1)$ **is a subset of** $s$**:**
As shown in Figure S18, a potential exon end site $e_m$ appears inside of a subpath $s = z_{[a:b]}(o_a, \ldots, o_b)$.

$$\mathsf{P}(s, Z_{I(e_m)} = 1, Z_{I(e_m)+1} = 1) \tag{161}$$
$$= \mathsf{P}(s) \tag{162}$$
$$= \mathsf{P}(Z_1 = 0, s) + \mathsf{P}(Z_1 = 1, s) \tag{163}$$
$$= (1-\pi) f_{01}(1, F(s)) w(s) + \pi f_{11}(1, F(s)) w(s) \tag{164}$$
$$= \big[(1-\pi) f_{01}(1, F(s)) + \pi f_{11}(1, F(s))\big] \times w(s). \tag{165}$$

**case 4** $F(s) \le I(e_m) < L(s)$ **and** $(Z_{I(e_m)} = 1, Z_{I(e_m)+1} = 1)$ **is not a subset of** $s$**:**
This event is now allowed.

$$\mathsf{P}(s, Z_{I(e_m)} = 1, Z_{I(s_m)+1} = 1) = 0 \tag{166}$$

- $\mathsf{P}(Z^n_{I(e_m)} = 1, Z^n_{I(e_m)+1} = 0, S_n = s)$

**case 1** $I(e_m) < F(s) - 1$**:**
As shown in Figure S18, a potential exon end site $e_m$ appears left side of a subpath $s = z_{[a:b]}(o_a, \ldots, o_b)$.

$$\mathsf{P}(Z_{I(e_m)} = 1, Z_{I(e_m)+1} = 0, s) \tag{167}$$
$$= \mathsf{P}(Z_1 = 0, Z_{I(e_m)} = 1, Z_{I(e_m)+1} = 0, s) + \mathsf{P}(Z_1 = 1, Z_{I(e_m)} = 1, Z_{I(e_m)+1} = 0, s) \tag{168}$$
$$= (1-\pi) f_{01}(1, I(e_m)) q_m f_{01}(I(s_m)+1, F(s)) w(s) + \pi f_{11}(1, I(s_m)) q_m f_{01}(I(s_m)+1, F(s)) w(s) \tag{169}$$
$$= \big[(1-\pi) f_{01}(1, I(e_m)) + \pi f_{11}(1, I(s_m))\big] \times q_m f_{01}(I(s_m)+1, F(s)) w(s). \tag{170}$$

**case 2** $L(s) \le I(e_m)$**:**
As shown in Figure S18, a potential exon end site $e_m$ appears right side of a subpath $s = z_{[a:b]}(o_a, \ldots, o_b)$.

$$\mathsf{P}(s, Z_{I(e_m)} = 1, Z_{I(e_m)+1} = 0) \tag{171}$$
$$= \mathsf{P}(Z_1 = 0, s, Z_{I(e_m)} = 1, Z_{I(e_m)+1} = 0) + \mathsf{P}(Z_1 = 1, s, Z_{I(e_m)} = 1, Z_{I(e_m)+1} = 0) \tag{172}$$
$$= (1-\pi) f_{01}(1, F(s)) w(s) f_{11}(L(s), I(s_m)) q_m + \pi f_{11}(1, F(s)) w(s) f_{11}(L(s), I(s_m)) q_m \tag{173}$$
$$= \big[(1-\pi) f_{01}(1, F(s)) + \pi f_{11}(1, F(s))\big] \times w(s) f_{11}(L(s), I(s_m)) q_m. \tag{174}$$

**case 3** $F(s) \leq I(e_m) < L(s)$ **and** $(Z_{I(e_m)} = 1, Z_{I(e_m)+1} = 0)$ **is a subset of** $s$**:**
As shown in Figure S18, a potential exon end site $e_m$ appears inside of a subpath $s = z_{[a:b]}(o_a, \ldots, o_b)$.

$$\begin{aligned}
&\mathsf{P}(s, Z_{I(e_m)} = 1, Z_{I(e_m)+0} = 1) && (175)\\
&= \mathsf{P}(s) && (176)\\
&= (1-\pi) f_{01}(1, F(s)) w(s) + \pi f_{11}(1, F(s)) w(s) && (177)\\
&= \big[(1-\pi) f_{01}(1, F(s)) + \pi f_{11}(1, F(s))\big] \times w(s). && (178)
\end{aligned}$$

**case 4** $F(s) \leq I(e_m) < L(s)$ **and** $(Z_{I(e_m)} = 1, Z_{I(e_m)+1} = 0)$ **is not a subset of** $s$**:**
This event is now allowed.

$$\mathsf{P}(s, Z_{I(e_m)} = 1, Z_{I(e_m)+1} = 0) = 0 \tag{179}$$

**case 5** $I(e_m) = F(s) - 1$**:**
This event is now allowed.

$$\mathsf{P}(s, Z_{I(e_m)} = 1, Z_{I(e_m)+1} = 0) = 0. \tag{180}$$

###### 2.4.1.7 $\mathsf{P}(Z^n_{I(s_m)} = 0, Z^n_{I(s_m)+1} = 1, s_l \in S_n, s_r \in S_n)$, $\mathsf{P}(Z^n_{I(s_m)} = 0, Z^n_{I(s_m)+1} = 0, s_l \in S_n, s_r \in S_n)$, $\mathsf{P}(Z^n_{I(e_m)} = 1, Z^n_{I(e_m)+1} = 0, s_l \in S_n, s_r \in S_n)$, and $\mathsf{P}(Z^n_{I(e_m)} = 1, Z^n_{I(e_m)+1} = 1, s_l \in S_n, s_r \in S_n)$

These probabilities can be derived analogously to Sections 2.4.1.6. Case 1 and case 2 in the paired-end setting apply whenever case 1 and case 2 apply in the single-end scenario for $s = s_l$ and $s = s_r$, respectively. Cases 3-5 need to consider if a start or end site is contained in $s_l$ as well as if it is contained in $s_r$. Additionally, the splice site might lie between the two mates $s_l$ and $s_r$.

- $\mathsf{P}(Z^n_{I(s_m)} = 0, Z^n_{I(s_m)+1} = 1, s_l \in S_n, s_r \in S_n)$
  **case 1** $I(s_m) < F(s_l)$**:**

$$\begin{aligned}
&\mathsf{P}(Z_{I(s_m)} = 0, Z_{I(s_m)+1} = 1, s_l, s_r) && (181)\\
&= \big[(1-\pi) f_{00}(1, I(s_m)) + \pi f_{10}(1, I(s_m))\big] \times p_m f_{11}(I(s_m)+1, F(s_l)) w_p((s_l, s_r). && (182)
\end{aligned}$$

  Here, $f_{**}(i, j)$ and $w_p(s_l, s_r)$ can be computed as described in Sections 2.4.1.3 and 2.4.1.5, respectively.

  **case 2** $L(s_r) < I(s_m)$**:**

$$\begin{aligned}
&\mathsf{P}(s_l, s_r, Z_{I(s_m)} = 0, Z_{I(s_m)+1} = 1) && (183)\\
&= \big[(1-\pi) f_{01}(1, F(s_l)) + \pi f_{11}(1, F(s_l))\big] \times w_p(s_l, s_r) f_{10}(L(s_r), I(s_m)) p_m. && (184)
\end{aligned}$$

  **case 3** $F(s_l) \leq I(s_m) < L(s_l)$ **and** $(Z_{I(s_m)} = 0, Z_{I(s_m)+1} = 1)$ **is a subset of** $s_l$**, or** $F(s_r) \leq I(s_m) < L(s_r)$ **and** $(Z_{I(s_m)} = 0, Z_{I(s_m)+1} = 1)$ **is a subset of** $s_r$**:**

$$\begin{aligned}
&\mathsf{P}(s_l, s_r, Z_{I(s_m)} = 0, Z_{I(s_m)+1} = 1) && (185)\\
&= \big[(1-\pi) f_{01}(1, F(s_l)) + \pi f_{11}(1, F(s_l))\big] \times w_p(s_l, s_r). && (186)
\end{aligned}$$

**case 4** $F(s_l) \leq I(s_m) < L(s_l)$ **and** $(Z_{I(s_m)} = 0, Z_{I(s_m)+1} = 1)$ **is not a subset of** $s_l$**, or** $F(s_r) \leq I(s_m) < L(s_r)$ **and** $(Z_{I(s_m)} = 0, Z_{I(s_m)+1} = 1)$ **is not a subset of** $s_r$**:**
This event is now allowed.

$$\mathsf{P}(s_l, s_r, Z_{I(s_m)} = 0, Z_{I(s_m)+1} = 1) = 0 \tag{187}$$

**case 5** $I(s_m) = L(s_r)$ **or** $I(s_m) = L(s_l)$**:**
This event is now allowed.

$$\mathsf{P}(s_l, s_r, Z_{I(s_m)} = 0, Z_{I(s_m)+1} = 1) = 0 \tag{188}$$

**case 6** $L(s_l) < I(s_m) < F(s_r)$**:**

$$\mathsf{P}(s_l, s_r, Z_{I(s_m)} = 0, Z_{I(s_m)+1} = 1) \tag{189}$$
$$= \big[(1-\pi) f_{01}(1, F(s_l)) + \pi f_{11}(1, F(s_l))\big] \times w'_p(s_l, s_r, z_{[I(s_m):I(s_m)+1]}(0,1)). \tag{190}$$

- $\mathsf{P}(Z^n_{I(s_m)} = 0, Z^n_{I(s_m)+1} = 0, s_l \in S_n, s_r \in S_n)$
  **case 1** $I(s_m) < F(s_l) - 1$**:**

$$\mathsf{P}(Z_{I(s_m)} = 0, Z_{I(s_m)+1} = 0, s_l, s_r) \tag{191}$$
$$= \big[(1-\pi) f_{00}(1, I(s_m)) + \pi f_{10}(1, I(s_m))\big] \times (1-p_m) f_{01}(I(s_m)+1, F(s_l)) w_p(s_l, s_r). \tag{192}$$

**case 2** $L(s_r) < I(s_m)$**:**

$$\mathsf{P}(s_l, s_r, Z_{I(s_m)} = 0, Z_{I(s_m)+1} = 0) \tag{193}$$
$$= \big[(1-\pi) f_{01}(1, F(s_l)) + \pi f_{11}(1, F(s_l))\big] \times w_p(s_l, s_r) f_{10}(L(s_r), I(s_m))(1-p_m). \tag{194}$$

**case 3** $F(s_l) \leq I(s_m) < L(s_l)$ **and** $(Z_{I(s_m)} = 0, Z_{I(s_m)+1} = 0)$ **is a subset of** $s_l$ **or** $F(s_r) \leq I(s_m) < L(s_r)$ **and** $(Z_{I(s_m)} = 0, Z_{I(s_m)+1} = 0)$ **is a subset of** $s_r$**:**

$$\mathsf{P}(s_l, s_r, Z_{I(s_m)} = 0, Z_{I(s_m)+0} = 1) \tag{195}$$
$$= \big[(1-\pi) f_{01}(1, F(s_l)) + \pi f_{11}(1, F(s_l))\big] \times w_p(s_l, s_r). \tag{196}$$

**case 4** $F(s_l) \leq I(s_m) < L(s_l)$ **and** $(Z_{I(s_m)} = 0, Z_{I(s_m)+1} = 0)$ **is not a subset of** $s_l$**, or** $F(s_r) \leq I(s_m) < L(s_r)$ **and** $(Z_{I(s_m)} = 0, Z_{I(s_m)+1} = 0)$ **is not a subset of** $s_r$**:**
This event is now allowed.

$$\mathsf{P}(s_l, s_r, Z_{I(s_m)} = 0, Z_{I(s_m)+1} = 0) = 0 \tag{197}$$

**case 5** $I(s_m) = F(s_l) - 1$ **or** $I(s_m) = L(s_l)$ **or** $I(s_m) = F(s_r) - 1$ **or** $I(s_m) = L(s_r)$**:**
This event is now allowed.

$$\mathsf{P}(s_l, s_r, Z_{I(s_m)} = 0, Z_{I(s_m)+1} = 0) = 0 \tag{198}$$

**case 6** $L(s_l) < I(s_m) < F(s_r) - 1$**:**

$$\mathsf{P}(s_l, s_r, Z_{I(s_m)} = 0, Z_{I(s_m)+0} = 1) \tag{199}$$
$$= \left[(1-\pi)f_{01}(1, F(s_l)) + \pi f_{11}(1, F(s_l))\right] \times w_p(s_l, s_r, z_{[I(s_m), I(s_m)+1]}(0,0)). \tag{200}$$

- $\mathsf{P}(Z^n_{I(e_m)} = 1, Z^n_{I(e_m)+1} = 1, s_l \in S_n, s_r \in S_n)$
  **case 1** $I(e_m) < F(s_l)$**:**

$$\mathsf{P}(Z_{I(e_m)} = 1, Z_{I(e_m)+1} = 1, s_l, s_r) \tag{201}$$
$$= \left[(1-\pi)f_{01}(1, I(e_m)) + \pi f_{11}(1, I(e_m))\right] \times (1-q_m) f_{11}(I(e_m)+1, F(s_l)) w_p(s_l, s_r). \tag{202}$$

**case 2** $L(s_r) \leq I(s_m)$**:**

$$\mathsf{P}(s_l, s_r, Z_{I(e_m)} = 1, Z_{I(e_m)+1} = 1) \tag{203}$$
$$= \left[(1-\pi)f_{01}(1, F(s_l)) + \pi f_{11}(1, F(s_l))\right] \times w_p(s_l, s_r) f_{11}(L(s_r), I(e_m))(1-q_m). \tag{204}$$

**case 3** $F(s_l) \leq I(e_m) < L(s_l)$ **and** $(Z_{I(e_m)} = 1, Z_{I(e_m)+1} = 1)$ **is a subset of** $s_l$**, or** $F(s_r) \leq I(e_m) < L(s_r)$ **and** $(Z_{I(e_m)} = 1, Z_{I(e_m)+1} = 1)$ **is a subset of** $s_r$**:**

$$\mathsf{P}(s_l, s_r, Z_{I(e_m)} = 1, Z_{I(e_m)+1} = 1) \tag{205}$$
$$= \left[(1-\pi)f_{01}(1, F(s_l)) + \pi f_{11}(1, F(s_l))\right] \times w_p(s_l, s_r). \tag{206}$$

**case 4** $F(s_l) \leq I(e_m) < L(s_l)$ **and** $(Z_{I(e_m)} = 1, Z_{I(e_m)+1} = 1)$ **is not a subset of** $s_l$**, or** $F(s_r) \leq I(e_m) < L(s_r)$ **and** $(Z_{I(e_m)} = 1, Z_{I(e_m)+1} = 1)$ **is not a subset of** $s_r$**:**
This event is now allowed.

$$\mathsf{P}(s_l, s_r, Z_{I(e_m)} = 1, Z_{I(s_m)+1} = 1) = 0 \tag{207}$$

**case 5** $L(s_l) \leq I(e_m) < F(s_r)$**:**

$$\mathsf{P}(s_l, s_r, Z_{I(e_m)} = 1, Z_{I(e_m)+1} = 1) \tag{208}$$
$$= \left[(1-\pi)f_{01}(1, F(s_l)) + \pi f_{11}(1, F(s_l))\right] \times w_p(s_l, s_r, z_{[I(e_m), I(e_m)+1]}(1,1)). \tag{209}$$

- $\mathsf{P}(Z^n_{I(e_m)} = 1, Z^n_{I(e_m)+1} = 0, s_l \in S_n, s_r \in S_n)$
  **case 1** $I(e_m) < F(s_l) - 1$**:**

$$\mathsf{P}(Z_{I(e_m)} = 1, Z_{I(e_m)+1} = 0, s_l, s_r) \tag{210}$$
$$= \left[(1-\pi)f_{01}(1, I(e_m)) + \pi f_{11}(1, I(s_m))\right] \times q_m f_{01}(I(s_m)+1, F(s_l)) w_p(s_l, s_r). \tag{211}$$

**case 2 $L(s_r) \leq I(e_m)$:**

$$\mathsf{P}(s_l, s_r, Z_{I(e_m)} = 1, Z_{I(e_m)+1} = 0) \tag{212}$$
$$= \big[(1-\pi) f_{01}(1, F(s_l)) + \pi f_{11}(1, F(s_l))\big] \times w_p(s_l, s_r) f_{11}(L(s_r), I(s_m)) q_m. \tag{213}$$

**case 3 $F(s_l) \leq I(e_m) < L(s_l)$ and $(Z_{I(e_m)} = 1, Z_{I(e_m)+1} = 0)$ is a subset of $s_l$, or $F(s_r) \leq I(e_m) < L(s_r)$ and $(Z_{I(e_m)} = 1, Z_{I(e_m)+1} = 0)$ is a subset of $s_r$:**

$$\mathsf{P}(s_l, s_r, Z_{I(e_m)} = 1, Z_{I(e_m)+0} = 1) \tag{214}$$
$$= \big[(1-\pi) f_{01}(1, F(s_l)) + \pi f_{11}(1, F(s_l))\big] \times w_p(s_l, s_r). \tag{215}$$

**case 4 $F(s_l) \leq I(e_m) < L(s_l)$ and $(Z_{I(e_m)} = 1, Z_{I(e_m)+1} = 0)$ is not a subset of $s_l$, or $F(s_r) \leq I(e_m) < L(s_r)$ and $(Z_{I(e_m)} = 1, Z_{I(e_m)+1} = 0)$ is not a subset of $s_r$:**
This event is now allowed.

$$\mathsf{P}(s_l, s_r, Z_{I(e_m)} = 1, Z_{I(e_m)+1} = 0) = 0 \tag{216}$$

**case 5 $I(s_m) = F(s_l) - 1$ or $I(s_m) = F(s_r) - 1$:**
This event is now allowed.

$$\mathsf{P}(s_l, s_r, Z_{I(e_m)} = 1, Z_{I(e_m)+1} = 0) = 0. \tag{217}$$

**case 6 $L(s_l) \leq I(e_m) < F(s_r) - 1$:**

$$\mathsf{P}(s_l, s_r, Z_{I(e_m)} = 1, Z_{I(e_m)+0} = 1) \tag{218}$$
$$= \big[(1-\pi) f_{01}(1, F(s_l)) + \pi f_{11}(1, F(s_l))\big] \times w_p(s_l, s_r, z_{[I(e_m), I(e_m)+1]}(1, 0)). \tag{219}$$

##### 2.4.2 EM algorithm

Let $Z^n = (Z_1^n, \ldots, Z_M^n)$ represent the isoform $T_n$, that is, the path from which read $n$ was derived. Then, the complete data likelihood, $\mathsf{P}(r,Z|\Theta) = \prod_{n=1}^{N} \mathsf{P}(r_n, Z^n|\Theta)$, can be written as

$$\prod_{n=1}^{N} \Big[ \sum_{(s,b):s\subset Z^n} \mathsf{P}(r_n, s_n = s, b_n = b, Z^n|\Theta) \Big] \tag{220}$$

where $s \subset Z^n$ means $s$ is a subpath of the path $Z^n$, (221)

$$= \prod_{n=1}^{N} \Big[ \sum_{(s,b):s\subset Z^n} \mathsf{P}(r_n|s_n = s, b_n = b)\mathsf{P}(s_n = s, b_n = b|Z^n)\mathsf{P}(Z^n|\Theta) \Big] \tag{222}$$

$$= \prod_{n=1}^{N} \Big[ \sum_{(s,b):s\subset Z^n,(s,b)\to r_n} 1 \frac{1}{l(Z^n)} \frac{l(Z^n) w_\Theta(Z^n)}{D(\Theta)} \Big] \tag{223}$$

$$= \prod_{n=1}^{N} \Big[ \sum_{(s,b):s\subset Z^n,(s,b)\to r_n} \frac{w_\Theta(Z^n)}{D(\Theta)} \Big] \tag{224}$$

$$= \prod_{n=1}^{N} \Big[ \frac{C(r_n, Z^n) w_\Theta(Z^n)}{D(\Theta)} \Big] \tag{225}$$

where $C(r_n, Z^n)$ indicates the number of $(s,b)$ in the isoform $T_n$ (defined by $Z^n$) which are matched to $r_n$. Then, we can rewrite it as

$$\begin{aligned} &\frac{1}{D(\Theta)^N} \prod_{n=1}^{N} \Big[ C(r_n, Z^n) \big[ \pi^{Z_1^n} (1-\pi)^{1-Z_1^n} \big] \\ &\times \Big[ \prod_{m=1}^{M_s} p_m^{(1-Z^n_{I(s_m)})(Z^n_{I(s_m)+1})} (1-p_m)^{(1-Z^n_{I(s_m)})(1-Z^n_{I(s_m)+1})} \Big] \\ &\times \Big[ \prod_{m=1}^{M_e} q_m^{(Z^n_{I(e_m)})(1-Z^n_{I(e_m)+1})} (1-q_m)^{(Z^n_{I(e_m)})(Z^n_{I(e_m)+1})} \Big] \Big]. \end{aligned} \tag{226}$$

And we can write a log likelihood $\log \mathsf{P}(r,Z|\Theta)$ as

$$-N \log D(\Theta) + \sum_{n=1}^{N} \log C(r_n, Z^n) + \sum_{n=1}^{N} Z_1^n \log \pi + \sum_{n=1}^{N} (1-Z_1^n) \log(1-\pi) \tag{227}$$

$$+ \sum_{n=1}^{N} \sum_{m=1}^{M_s} \Big[ (1-Z^n_{I(s_m)})(Z^n_{I(s_m)+1}) \log p_m \Big] + \sum_{n=1}^{N} \sum_{m=1}^{M_s} \Big[ (1-Z^n_{I(s_m)})(1-Z^n_{I(s_m)+1}) \log(1-p_m) \Big] \tag{228}$$

$$+ \sum_{n=1}^{N} \sum_{m=1}^{M_e} \Big[ (Z^n_{I(e_m)})(1-Z^n_{I(e_m)+1}) \log q_m \Big] + \sum_{n=1}^{N} \sum_{m=1}^{M_e} \Big[ (Z^n_{I(e_m)})(Z^n_{I(e_m)+1}) \log(1-q_m) \Big]. \tag{229}$$

Note that the transition probabilities in our model don't allow isoforms where $Z_{I(s_m)} = 1$ and $Z_{I(s_m)+1} = 0$ at any exon start site and $Z_{I(e_m)} = 0$ and $Z_{I(e_m)+1} = 1$ at any exon end site, and $C(r_n, Z^n)$ does not depend on $\Theta$.

###### 2.4.2.1 E-step

We denote by $\Theta^l$ the model parameters at step $l$ of the EM algorithm.

$$\mathsf{P}((1-Z^n_{I(s_m)})(Z^n_{I(s_m)+1}) = 1|r_n, \Theta^l) = \frac{\mathsf{P}((1-Z^n_{I(s_m)})(Z^n_{I(s_m)+1}) = 1, r_n, \Theta^l)}{\mathsf{P}(r_n, \Theta^l)} \tag{230}$$

$$= \frac{\mathsf{P}(Z^n_{I(s_m)} = 0, Z^n_{I(s_m)+1} = 1, S_n = s, \Theta^l)}{\mathsf{P}(S_n = s, \Theta^l)}, \tag{231}$$

where $s$ is the shortest subpath from which read $n$ is derived, and $\mathsf{P}(Z^n_{I(s_m)} = 0, Z^n_{I(s_m)+1} = 1, S_n = s, \Theta^l)$ is given in Section 2.4.1.6. And

$$\mathsf{P}(S_n = s, \Theta^l) \tag{232}$$

$$= \mathsf{P}(Z_1 = 0, S_n = s, \Theta^l) + \mathsf{P}(Z_1 = 1, S_n = s, \Theta^l) \tag{233}$$

$$= \mathsf{P}(Z_1 = 0)\mathsf{P}(Z_{F(s)} = 1|Z_1 = 0)\mathsf{P}(S_n = s|Z_{F(s)} = 1) + \mathsf{P}(Z_1 = 1)\mathsf{P}(Z_{F(s)} = 1|Z_1 = 1)\mathsf{P}(S_n = s|Z_{F(s)} = 1) \tag{234}$$

$$= (1-\pi)f_{01}(1, F(s))w(s) + \pi f_{11}(1, F(s))w(s), \tag{235}$$

where $f_{**}(i,j)$ and $w(s)$ are given in Sections 2.4.1.3 and 2.4.1.4.

We note that reads mapping to the same signature have the same subpath for $S_n$ (i.e., the shortest subpath of $T_n$ from which read $n$ is derived), so they have the same posterior probability of $(1-Z^n_{I(s_m)})(Z^n_{I(s_m)+1}) = 1$. That is, for $r_n$ and $r_{n'}$ mapping to the same signature, $\mathsf{P}((1-Z^n_{I(s_m)})(Z^n_{I(s_m)+1}) = 1|r_n, \Theta^l) = \mathsf{P}((1-Z^{n'}_{I(s_m)})(Z^{n'}_{I(s_m)+1}) = 1|r_{n'}, \Theta^l)$.

Similarly,

$$\mathsf{P}((1-Z^n_{I(s_m)})(1-Z^n_{I(s_m)+1}) = 1|r_n, \Theta^l) = \frac{\mathsf{P}((1-Z^n_{I(s_m)})(1-Z^n_{I(s_m)+1}) = 1, r_n, \Theta^l)}{\mathsf{P}(r_n, \Theta^l)} \tag{236}$$

$$= \frac{\mathsf{P}(Z^n_{I(s_m)} = 0, Z^n_{I(s_m)+1} = 0, S_n = s, \Theta^l)}{\mathsf{P}(S_n = s, \Theta^l)}, \tag{237}$$

$$\mathsf{P}((Z^n_{I(e_m)})(1-Z^n_{I(e_m)+1}) = 1|r_n, \Theta^l) = \frac{\mathsf{P}((Z^n_{I(e_m)})(1-Z^n_{I(e_m)+1}) = 1, r_n, \Theta^l)}{\mathsf{P}(r_n, \Theta^l)} \tag{238}$$

$$= \frac{\mathsf{P}(Z^n_{I(e_m)} = 1, Z^n_{I(e_m)+1} = 0, S_n = s, \Theta^l)}{\mathsf{P}(S_n = s, \Theta^l)}, \tag{239}$$

$$\mathsf{P}((Z^n_{I(e_m)})(Z^n_{I(e_m)+1}) = 1|r_n, \Theta^l) = \frac{\mathsf{P}((Z^n_{I(e_m)})(Z^n_{I(e_m)+1}) = 1, r_n, \Theta^l)}{\mathsf{P}(r_n, \Theta^l)} \tag{240}$$

$$= \frac{\mathsf{P}(Z^n_{I(e_m)} = 1, Z^n_{I(e_m)+1} = 1, S_n = s, \Theta^l)}{\mathsf{P}(S_n = s, \Theta^l)}, \tag{241}$$

where $\mathsf{P}(Z^n_{I(s_m)}=0, Z^n_{I(s_m)+1}=0, S_n=s, \Theta^l)$, $\mathsf{P}(Z^n_{I(e_m)}=1, Z^n_{I(e_m)+1}=0, S_n=s, \Theta^l)$, $\mathsf{P}(Z^n_{I(e_m)}=1, Z^n_{I(e_m)+1}=1, S_n=s, \Theta^l)$ are given in Section 2.4.1.6. Also, we note that reads mapping to the same signature have the same posterior probabilities.

###### 2.4.2.2 M-step

We find the parameters $\Theta=(\pi, p_1, \dots, p_{M_s}, q_1, \dots, q_{M_e})$ which maximizes $\mathrm{E}_{Z|r,\Theta^l}[\log\mathsf{P}(r,Z|\Theta)]$.

$$\Theta^{l+1} = \underset{\Theta}{\operatorname{argmax}} \, \mathrm{E}_{Z|r,\Theta^l}[\log \mathsf{P}(r, Z \mid \Theta)] \tag{242}$$

$$\equiv \underset{\Theta}{\operatorname{argmax}} \, Q(\Theta \mid \Theta^l). \tag{243}$$

In the Section 2.4.2, we showed that $\log\mathsf{P}(r,Z|\Theta)=\log l(\Theta)$ can be written as

$$-N\log D(\Theta) + \sum_{n=1}^{N}\log C(r_n, Z^n) + \sum_{n=1}^{N} Z_1^n \log\pi + \sum_{n=1}^{N}(1-Z_1^n)\log(1-\pi) \tag{244}$$

$$+\sum_{n=1}^{N}\sum_{m=1}^{M_s}\left[(1-Z^n_{I(s_m)})(Z^n_{I(s_m)+1})\log p_m\right] + \sum_{n=1}^{N}\sum_{m=1}^{M_s}\left[(1-Z^n_{I(s_m)})(1-Z^n_{I(s_m)+1})\log(1-p_m)\right] \tag{245}$$

$$+\sum_{n=1}^{N}\sum_{m=1}^{M_e}\left[(Z^n_{I(e_m)})(1-Z^n_{I(e_m)+1})\log q_m\right] + \sum_{n=1}^{N}\sum_{m=1}^{M_e}\left[(Z^n_{I(e_m)})(Z^n_{I(e_m)+1})\log(1-q_m)\right]. \tag{246}$$

Note that $C(r_n, Z^n)$ does not depend on $\Theta$.

- **$p_m^{l+1}$ for $m=1,\dots,M_s$**
  Let $p'_m = 1-p_m$. Then,

$$\frac{\partial}{\partial p_m}Q(\Theta\mid\Theta^l) = -N\frac{1}{D(\Theta)}\frac{\partial D(\Theta)}{\partial p_m} + \frac{1}{p_m}\sum_{n=1}^{N}\mathsf{P}((1-Z^n_{I(s_m)})(Z^n_{I(s_m)+1})=1|r_n,\Theta^l)=0, \tag{247}$$

$$\frac{\partial}{\partial p'_m}Q(\Theta\mid\Theta^l) = -N\frac{1}{D(\Theta)}\frac{\partial D(\Theta)}{\partial p'_m} + \frac{1}{p'_m}\sum_{n=1}^{N}\mathsf{P}((1-Z^n_{I(s_m)})(1-Z^n_{I(s_m)+1})=1|r_n,\Theta^l)=0, \tag{248}$$

  leading to

$$p_m = \frac{\sum_{n=1}^{N}\mathsf{P}((1-Z^n_{I(s_m)})(Z^n_{I(s_m)+1})=1|r_n,\Theta^l)}{\frac{N}{D(\Theta)}\frac{\partial D(\Theta)}{\partial p_m}}, \tag{249}$$

$$p'_m = \frac{\sum_{n=1}^{N}\mathsf{P}((1-Z^n_{I(s_m)})(1-Z^n_{I(s_m)+1})=1|r_n,\Theta^l)}{\frac{N}{D(\Theta)}\frac{\partial D(\Theta)}{\partial p'_m}}. \tag{250}$$

  We compute $\frac{\partial D(\Theta)}{\partial p_m}$ and $\frac{\partial D(\Theta)}{\partial p'_m}$ as follows. $D(\Theta)$ can be written

$$D(\Theta) = E(l(T)) = E(l(Z_{[1:M]})) \tag{251}$$

$$= E(l(Z_{[1:M]}), Z_{I(S_m)}=0, Z_{I(S_m)+1}=1) + E(l(Z_{[1:M]}), Z_{I(S_m)}=0, Z_{I(S_m)+1}=0) \tag{252}$$

$$+ E(l(Z_{[1:M]}), Z_{I(S_m)}=1, Z_{I(S_m)+1}=0) + E(l(Z_{[1:M]}), Z_{I(S_m)}=1, Z_{I(S_m)+1}=1). \tag{253}$$

Because $p_m = \mathsf{P}(Z_{I(S_m)+1} = 1|Z_{I(S_m)} = 0)$ and $p'_m = \mathsf{P}(Z_{I(S_m)+1} = 0|Z_{I(S_m)} = 0)$, the last two components do not contain $p_m$ or $p'_m$ and the first two components can be re-written as

$$\mathsf{P}(Z_{I(S_m)} = 0)p_m[E(l(Z_{[1:I(S_m)]})|Z_{I(S_m)} = 0, Z_{I(S_m)+1} = 1) + E(l(Z_{[I(S_m)+1:M]})|Z_{I(S_m)} = 0, Z_{I(S_m)+1} = 1)] \tag{254}$$

$$+ \mathsf{P}(Z_{I(S_m)} = 0)p'_m[E(l(Z_{[1:I(S_m)]})|Z_{I(S_m)} = 0, Z_{I(S_m)+1} = 0) + E(l(Z_{[I(S_m)+1:M]})|Z_{I(S_m)} = 0, Z_{I(S_m)+1} = 0)] \tag{255}$$

$$= \mathsf{P}(Z_{I(S_m)} = 0)p_m[E(l(Z_{[1:I(S_m)]})|Z_{I(S_m)} = 0) + E(l(Z_{[I(S_m)+1:M]})|Z_{I(S_m)+1} = 1)] \tag{256}$$

$$+ \mathsf{P}(Z_{I(S_m)} = 0)p'_m[E(l(Z_{[1:I(S_m)]})|Z_{I(S_m)} = 0) + E(l(Z_{[I(S_m)+1:M]})|Z_{I(S_m)+1} = 0)] \tag{257}$$

$$= \mathsf{P}(Z_{I(S_m)} = 0)\left(p_m[l_p(I(S_m), \text{out}) + l_s(I(S_m)+1, \text{in})] + p'_m[l_p(I(S_m), \text{out}) + l_s(I(S_m)+1, \text{out})]\right) \tag{258}$$

Then,

$$\frac{\partial D(\Theta)}{\partial p_m} = \mathsf{P}(Z_{I(S_m)} = 0)[l_p(I(S_m), \text{out}) + l_s(I(S_m)+1, \text{in})], \tag{259}$$

$$\frac{\partial D(\Theta)}{\partial p'_m} = \mathsf{P}(Z_{I(S_m)} = 0)[l_p(I(S_m), \text{out}) + l_s(I(S_m)+1, \text{out})]. \tag{260}$$

Let

$$A_m = \frac{\sum_{n=1}^N \mathsf{P}((1 - Z^n_{I(s_m)})(Z^n_{I(s_m)+1}) = 1|r_n, \Theta^l)}{l_p(I(s_m), \text{out}) + l_s(I(s_m)+1, \text{in})}, \tag{261}$$

$$B_m = \frac{\sum_{n=1}^N \mathsf{P}((1 - Z^n_{I(s_m)})(1 - Z^n_{I(s_m)+1}) = 1|r_n, \Theta^l)}{l_p(I(s_m), \text{out}) + l_s(I(s_m)+1, \text{out})}. \tag{262}$$

Then,

$$p_m = \frac{A_m}{\frac{N}{D(\Theta)}\mathsf{P}(Z_{I(S_m)} = 0)}, \tag{263}$$

$$p'_m = \frac{B_m}{\frac{N}{D(\Theta)}\mathsf{P}(Z_{I(S_m)} = 0)}. \tag{264}$$

Because $p_m + p'_m = 1$,

$$\frac{A_m + B_m}{\frac{N}{D(\Theta)}\mathsf{P}(Z_{I(S_m)} = 0)} = 1, \tag{265}$$

resulting in

$$\frac{N}{D(\Theta)}\mathsf{P}(Z_{I(S_m)} = 0) = A_m + B_m. \tag{266}$$

Thus,

$$p_m^{l+1} = p_m = \frac{A_m}{A_m + B_m}. \tag{267}$$

Let $c = (c_j)_{j=1}^{J}$ represent the signature counts over $J$ signatures and $s_j$ for $j = 1, \ldots, J$ represent a path corresponding to the $j$-th signature. Then, $A_m$ and $B_m$ can be written as

$$A_m = \frac{\sum_{j=1}^{J} c_j \mathsf{P}((1 - Z^j_{I(s_m)})(Z^j_{I(s_m)+1}) = 1 | s_j, \Theta^l)}{l_p(I(s_m), \text{out}) + l_s(I(s_m)+1, \text{in})}, \tag{268}$$

$$B_m = \frac{\sum_{j=1}^{J} c_j \mathsf{P}((1 - Z^j_{I(s_m)})(1 - Z^j_{I(s_m)+1}) = 1 | s_j, \Theta^l)}{l_p(I(s_m), \text{out}) + l_s(I(s_m)+1, \text{out})}. \tag{269}$$

So $p_m^{l+1}$ can be computed only using the signature counts.

- **$q_m^{l+1}$ for $m = 1, \ldots, M_e$:**
  Using the derivation similar to one for $p_m^{l+1}$ above, we can obtain the following result.
  Let

$$C_m = \frac{\sum_{n=1}^{N} \mathsf{P}((Z^n_{I(e_m)})(1 - Z^n_{I(e_m)+1}) = 1 | r_n, \Theta^l)}{l_p(I(s_m), \text{in}) + l_s(I(s_m)+1, \text{out})}, \tag{270}$$

$$D_m = \frac{\sum_{n=1}^{N} \mathsf{P}((Z^n_{I(e_m)})(Z^n_{I(e_m)+1}) = 1 | r_n, \Theta^l)}{l_p(I(s_m), \text{in}) + l_s(I(s_m)+1, \text{in})}. \tag{271}$$

  Then

$$q_m^{l+1} = \frac{C_m}{C_m + D_m}. \tag{272}$$

  Using the derivation similar to one for $p_m^{l+1}$ above, we can show that $q_m^{l+1}$ can be computed only using the signature counts.

- $\pi^{l+1}$
  Using the derivation similar to one for $p_m^{l+1}$ above, we can obtain the following result.
  Let

$$E = \frac{\sum_{n=1}^{N} \mathsf{P}(Z^n_1 = 1 | r_n, \Theta^l)}{l_s(1, \text{in})}, \tag{273}$$

$$F = \frac{\sum_{n=1}^{N} \mathsf{P}((Z^n_1 = 0 | r_n, \Theta^l)}{l_s(1, \text{out})}. \tag{274}$$

  Then

$$\pi^{l+1} = \frac{E}{E + F}. \tag{275}$$

  Using the derivation similar to one for $p_m^{l+1}$ above, we can show that $\pi^{l+1}$ can be computed only using the signature counts.

#### 2.5 Tools and parameters

##### 2.5.1 Polyester simulator

We used simulated data to evaluate McSplicer accuracy. As mentioned in the main text, we used Polyester simulator (version 1.16.0) to simulate RNA-seq reads [3] from $Homo\_sapiens.GRCh38.91.cdna.fa$. For the three different sequencing depths, we used the software with its default parameters, and we ran it under the following environment:

```
R version 3.5.2 (2018-12-20)
Platform: x86_64-redhat-linux-gnu (64-bit)
Running under: Scientific Linux 7.5 (Nitrogen)
```

As previously mentioned, we provided Polyester with ground truth abundances computed by running RSEM quantification tool [5] on RNA-seq data obtained from short read archive, dataset ID: SRR6987574 [2]. Then, we randomly selected a set of 1000 genes which have at least two expressed transcripts and have high gene expression level, i.e., gene-level read count per kilobase $> 500$.

##### 2.5.2 STAR aligner

The simulated reads were aligned back to the human reference genome (GRCh38.91) by running STAR (version 2.5.4b) [1] with the following parameters:

```
--outSAMtype BAM SortedByCoordinate
--sjdbGTFfile Homo_sapiens.GRCh38.91.gtf
--runThreadN 8
--readFilesIn {ployester_output.fasta}
--outFileNamePrefix {output_prefix}
--genomeDir {genome_directory}
```

The remaining set of parameters were left to the default values.

For indexing the alignment BAM files (i.e., STAR output) we used Samtools (version 0.1.8) [6].

##### 2.5.3 StringTie

We ran StringTie [7] (version 1.3.4d) with a genome-guided mode enabled (-G option). The remaining parameters of StringTie were left to the default values.

##### 2.5.4 SplAdder

We ran SplAdder with the following set of parameters for benchmarking on simulated data:

```
--bams {bam_files}
--annotation {annotation_gtf}
--merge_strat merge_graphs
--event_types exon_skip,intron_retention,alt_3prime,
alt_5prime,mult_exon_skip
```

[2] http://www.ncbi.nlm.nih.gov/sra

```
--confidence 2
--pyproc n
--compress_text n
--ignore_mismatches
--outdir {output_directory}
```

We set the *confidence* parameter to 1 when running SplAdder on the SIRV dataset in order to detect novel events.

##### 2.5.5 MAJIQ

We ran MAJIQ (version 2.0) with the default parameters but with de novo option disabled, i.e., $disable-denovo$ for all the benchmark on simulated data experiments. The main reason behind that was when running MAJIQ without $disable-denovo$ parameter (i.e., enabling de novo mode), we noticed many false positive events. We enable de novo mode again when evaluating MAJIQ on SIRV dataset to detect as many novel events as possible.
